# Detecting cell segmentation errors using doublet methods

**DOI:** 10.64898/2026.09.15.751798

**Authors:** Ameer Sarwar, Jesse Gillis

## Abstract

In spatial transcriptomics, cell segmentation is used to draw boundaries around cells. Molecules located within a cell’s boundary are assigned to it, making its gene expression profile dependent on segmentation accuracy. To identify potentially problematic cells, studies increasingly use doublet detection methods, a class of algorithms developed to recognize molecular admixture from two cells in scRNA-seq. To evaluate their suitability in spatial data, we model cell segmentation errors as a continuum of partial molecular admixture between neighboring cells, generated by varying the loss of a cell’s own transcripts and the gain of transcripts from its neighbor. Evaluating 8 doublet methods across 16 spatial datasets, we characterize the conditions under which they perform well and identify their failure modes. Detection improves with increasing molecular admixture and transcriptional dissimilarity between neighboring cells, with cxds2, scDblFinder.score, and a simple baseline (unique_genes) performing best. These patterns are consistent across datasets spanning different gene panels, technological platforms, staining techniques, segmentation algorithms, and error frequencies. In unperturbed spatial data, elevated doublet scores localize to regions consistent with segmentation problems, suggesting that they can help prioritize cells or regions for inspection, segmentation refinement, or transcript reassignment. Our results establish when doublet methods can provide useful quality control signals for cell segmentation, supporting their use in spatial transcriptomics.

## Introduction

Imaging-based spatial transcriptomic technologies resolve individual RNA transcripts *in situ* and assign them to cells based on their physical locations^1,2^. Because these assays do not directly capture cell boundaries, the boundaries must be computationally inferred though cell segmentation^3–26^, typically using auxiliary imaging stains or transcript point clouds. Segmentation is a challenging problem because tissues contain densely packed cells with heterogeneous sizes and morphologies, closely apposed or overlapping structures, and variable image quality. Low-quality cell segmentation can distort cellular expression profiles and produce misleading biological conclusions^27–30^. Highlighting this problem, Mitchell et al.^30^ performed differential expression analysis on the same cell type in distinct spatial locations. They identified markers of neighboring cell types among the differentially expressed genes, consistent with transcript misassignment between adjacent cells. Despite rapid developments in segmentation algorithms^3–26^, poorly segmented cells typically remain in the data, requiring robust quality control methods to reduce the risk of incorrect biological conclusions.

Doublet detection methods^31–36^ are frequently used in spatial transcriptomics to identify poorly segmented cells^15/09^/2026 1:10:00 PM (**Fig. 1a**). These methods were originally developed for single-cell RNA sequencing (scRNA-seq), where they identify droplets containing two or more cells. Methods such as Scrublet^32^ and DoubletFinder^31^ score observed transcriptomes according to their similarity to *in silico*-generated doublets and can detect profiles containing contributions from transcriptionally distinct cell populations. Because segmentation errors can also produce molecular admixture^27– 30^, these methods have been widely applied to spatial transcriptomic data, including large-scale atlases that have subsequently supported numerous studies. Indeed, they are increasingly being incorporated into standard data processing pipelines^37–41^. However, it remains unclear whether the concept of a doublet – and the algorithms designed to detect them – can be directly applied to spatial transcriptomics.

**Figure 1.**
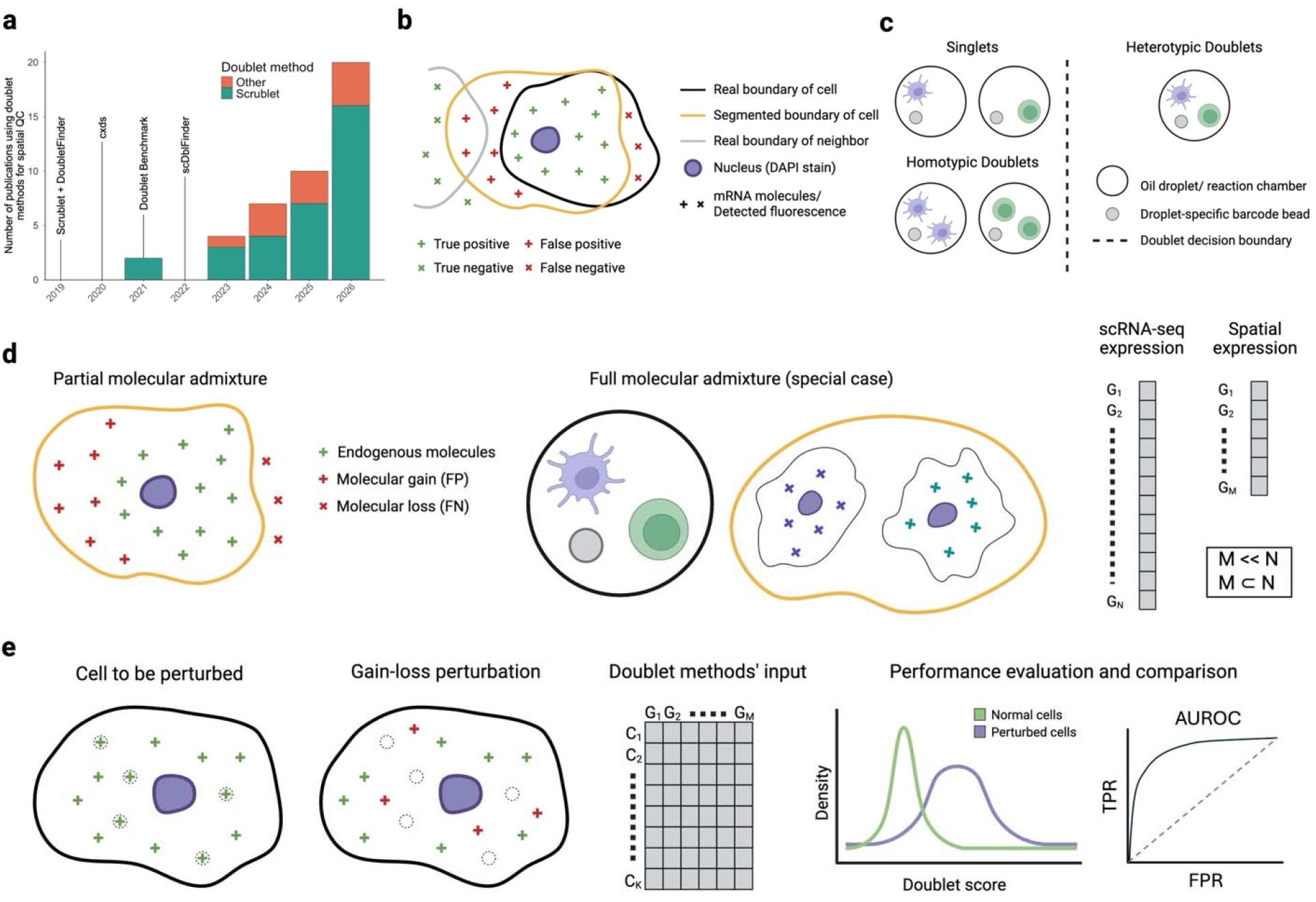
Cell segmentation error modeling and evaluation scheme. **a**, Studies using doublet detection methods for quality control in imaging-based spatial transcriptomics (total = 43). Annotations indicate publication dates of selected methods and a benchmarking study in scRNA-seq. Year 2026 includes studies until July. **b**, Segmentation errors can lead to misassigned transcripts, where the observed gene expression profile (yellow) contains external transcripts and misses internal transcripts. **c**, Doublet methods identify droplets containing two or more cells, which are usually transcriptionally different from each other (heterotypic). **d**, Left, segmentation errors can result in partial molecular admixture (see **b**). Middle, doublets form when two complete cells are co-encapsulated. In spatial transcriptomics, co-encapsulation occurs only when a segmentation boundary contains two cells. Right, scRNA-seq offers transcriptome-wide measurements but spatial technologies usually target 100-1000 genes. **e**, Cell segmentation errors are modeled as partial molecular admixture, where perturbed cells lose their transcripts and gain other cells’ transcripts. The resultant gene expression matrix is inputted into doublet methods, whose scores are used to calculate AUROC. Dashed gray line indicates random performance, AUROC = 0.5. Created in Biorender.com.

A doublet contains RNA from at least two cells and may therefore comprise substantial contributions from two transcriptomes^31–36^. By contrast, segmentation errors^27–30^ span a continuum: a cell boundary may capture only a small number of transcripts from a neighboring cell, exclude part of the target cell, or in more extreme cases, encompass multiple cells. Defining a “spatial doublet” only as a segmented region containing two complete cells therefore overlooks the partial and asymmetric molecular admixture characteristic of spatial data. Beyond this conceptual mismatch, technical differences between the two assay types further complicate the direct application of doublet detection methods to spatial data. Doublet methods were developed for the transcriptome-wide measurements generated by scRNA-seq^31–36^, whereas imaging-based spatial assays typically profile targeted panels of approximately 100-1,000 genes^1,2^. Differences in sample handling, capture chemistry, gene coverage, and data processing pipelines introduce further distributional shifts and batch effects between scRNA-seq and spatial transcriptomic data^42,43^. Together, these conceptual and technical differences raise uncertainty about whether conventional doublet detection methods can reliably identify segmentation errors in spatial transcriptomics.

Here, we model cell segmentation errors as partial molecular admixture, in which target cells gain transcripts from neighboring cells while losing their own transcripts. We benchmark eight doublet detection methods^31–36^ and two baseline approaches across 16 adult mouse brain datasets^44–59^ spanning seven technologies, multiple staining and segmentation strategies, and diverse gene panel designs. The mouse brain provides a challenging but well-characterized^60–62^ setting because of its morphological complexity and transcriptional heterogeneity. We find that cxds2, scDblFinder.score, and the number of uniquely expressed genes achieve the strongest discrimination of simulated segmentation errors, with AUROCs of approximately 0.8. Performance is driven primarily by the severity of admixture and the transcriptional dissimilarity between the target and contaminating cells, such that subtle errors between similar neighbors are particularly difficult to identify. Elevated doublet scores are associated with patterns consistent with segmentation issues, including dense transcript point clouds and limited separation between adjacent cells. Together, our work characterizes the conditions under which doublet detection methods are informative for spatial transcriptomic quality control and supports their use for prioritizing suspect cells for re-segmentation or transcript re-assignment^30,37–41,63–72^, while underscoring that they remain incomplete and context-dependent proxies for segmentation quality.

## Results

### Cell segmentation error as partial molecular admixture and doublet-detection evaluation scheme

Cell segmentation errors can lead to transcript misassignment. The segmented cell may contain transcripts from the true underlying cell and from the background, including neighboring cells or the extracellular space. At the same time, it can exclude transcripts belonging to the true cell. Thus, the observed expression profile might be a partial admixture of multiple sources, with the true profile being under-represented (**Fig. 1b**). By contrast, doublets in scRNA-seq contain two complete cells. To identify them, most doublet detection methods^31,32,36^ compare droplets’ expression profiles to those of *in silico*-generated doublets while other methods find ectopic co-expression patterns^33^. The central insight is that doublets exhibit admixed profiles that differ from the gene expression patterns of genuine cells. These methods excel at identifying heterotypic doublets (different cell types), while homotypic doublets (similar cell types) are harder to detect but are generally considered less consequential^34,35^ (**Fig. 1c**).

Segmentation errors pose a harder detection problem than doublets for two reasons. First, they span a continuum of admixture severity: a segmented cell might gain or lose anywhere from a small fraction to the majority of its transcripts (**Fig. 1d**, left). A doublet represents one fixed, extreme case: complete admixture with no loss (**Fig. 1d**, middle). Many segmentation errors are therefore far more subtle than the doublets these methods were designed to detect. Second, imaging-based spatial assays profile only 100-1,000 genes compared to the transcriptome-wide coverage of scRNA-seq (**Fig. 1d**, right), leaving far less signal from which to detect any given error. Doublet detection methods applied to spatial data must therefore identify more subtle admixture using less information, motivating a systematic evaluation of their generalizability and robustness.

To this end, we model cell segmentation errors as partial molecular admixture, where the cell loses a proportion of its transcripts and gains a proportion of another cell’s transcripts. We vary the gain and loss proportions independently in 10% increments, so that a cell might, for instance, gain 30% of another cell’s transcripts and lose 10% of its own transcripts. The resultant gene expression matrix contains perturbed and unperturbed cells and is the sole input to doublet detection methods. The methods’ per-cell doublet scores are used together with the perturbation status to calculate AUROC (**Fig. 1e**), thereby quantifying error-detection for a given perturbation.

### Doublet methods detect partial molecular admixture in scRNA-seq data

Single-cell RNA-seq captures complete cells without cell segmentation, making it an appropriate reference data for our study. Its broad gene coverage facilitates panel-specific experiments on the same set of cells. Across 16 spatial panels, the gene coverage ranges from 98 to 1122 (**Fig. 2a**). Most of these genes appear in 2-3 panels and very few are present in more than half (**Fig. S1a**, power-law distribution), with 60 genes shared between two panels on average (**Fig. S1b-c**). Our collection thus includes studies with diverse gene panels, enabling tests of doublet methods’ robustness to panel composition.

**Figure 2.**
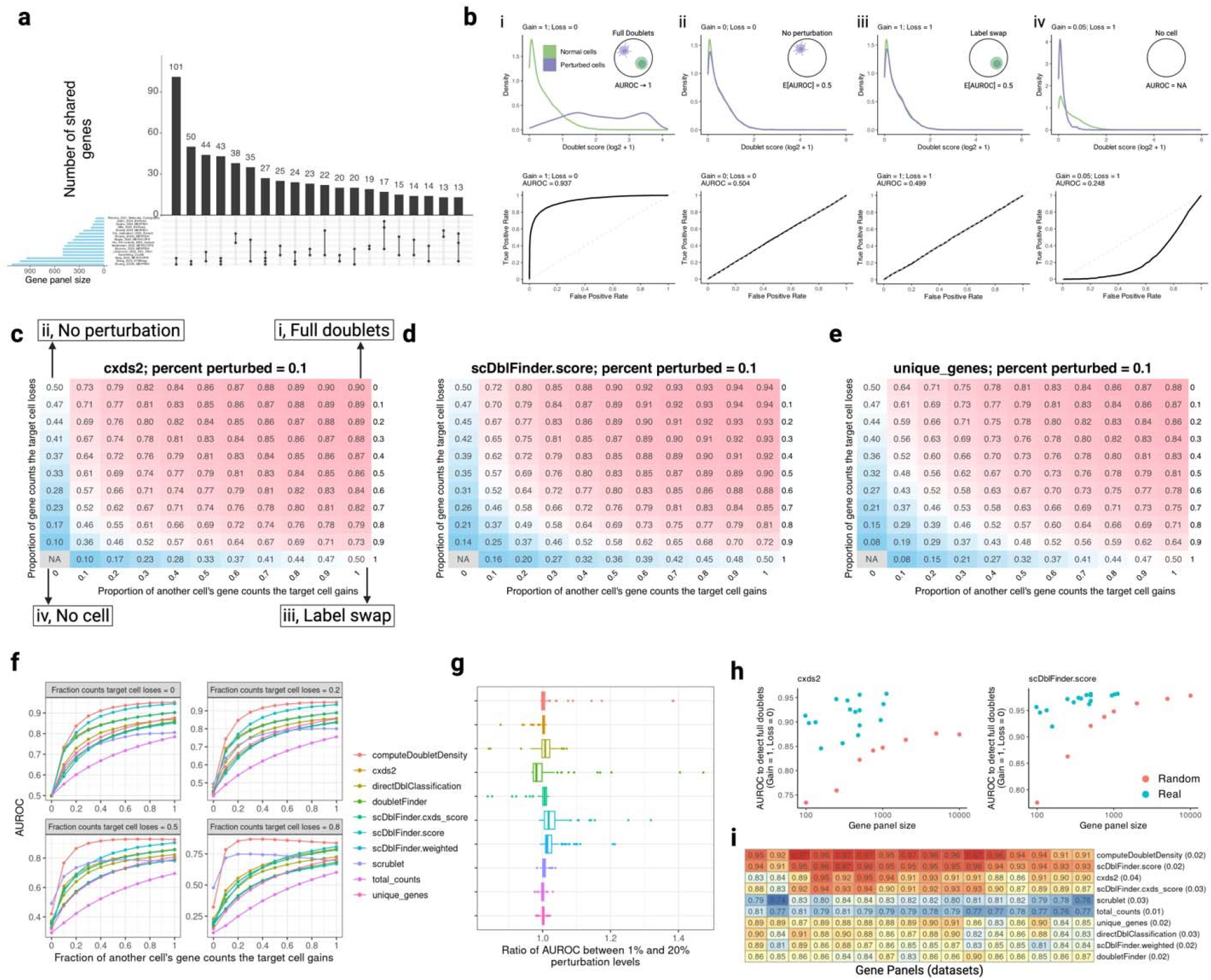
Error detection in scRNA-seq data. **a**, Upset plot shows overlaps between and sizes of 16 gene panels. **b**, Top, doublet score distributions colored by perturbation status and faceted by type (i-iv), with insets showing the associated perturbations. In iv, inset shows gain = 0 and loss = 1, but the distributions correspond to gain = 0.05 and loss = 1 (see text). All four plots show computeDoubletDensity scores on the Vizgen gene panel. Bottom, ROC curves correspond to perturbation type (i-iv) and AUROC quantifies the overall performance. **c-e**, Heatmaps show AUROCs across the perturbation space for cxds2 (**c**), scDblFinder.score (**d**), and unique_genes (**e**). Labels in (**c**) correspond to example perturbations in (**b**). Each entry shows AUROC averaged over 16 gene panels. In each case, 10% of cells were perturbed. **f**, AUROC against admixture severity, faceted by transcript loss. **g**, Ratio of AUROC when 1% and 20% of cells are perturbed. Ratios are taken between corresponding entries in the perturbation space. Legend is the same as in (**f**). **h**, AUROC (full doublets) against gene panel size for real and simulated (random) panels, with cxds2 and scDblFinder.score as examples. Data points for simulated panels are the average across 100 runs. **i**, AUROC (full doublets) of methods against gene panels. Numbers next to methods are standard deviations across panels. See also Figures S1-4. Created in Biorender.com.

To explain the relationship between perturbations and AUROC scores, we applied computeDoubletDensity to reference scRNA-seq data (Allen Institute adult mouse whole-brain atlas) filtered to the Vizgen MERSCOPE gene panel. Our sample contains ∼50,000 cells, matching the average number of cells in a spatial slice. A perturbation of gain = 1 and loss = 0 generates a full doublet: one complete cell is gained without loss of target cell. computeDoubletDensity here generates distinct doublet score distributions for perturbed and unperturbed cells (**Fig. 2b i**, AUROC = 0.937). A perturbation of gain = 0 and loss = 0 is equivalent to no perturbation while gain = 1 and loss = 1 replaces the target cell with the donor cell. In both cases, unperturbed and “perturbed” cells have similar doublet score distributions, as expected (**Fig. 2b ii-iii**, AUROC = 0.5). Gain = 0 and loss = 1 creates an empty expression vector, preventing methods’ execution, e.g., due to division by 0. Thus, we performed gain = 0.05 and loss = 1, which generates a cell fragment as the perturbed cell. Because fragments are subsets of genuine cells, they are seen as purer (less doublet-like) than real cells, generating *lower* doublet scores than unperturbed cells (**Fig. 2b iv**, AUROC = 0.248).

Independently varying gain and loss generates a perturbation *space*. After filtering scRNA-seq for a gene panel, we apply a gain-loss perturbation and input the data to doublet methods, whose scores are used to calculate AUROCs. We evaluated the methods across the perturbation space, where entries show AUROCs averaged over 16 panels. Looking at cxds2 (co-expression), scDblFinder.score (*in silico* doublets; best in benchmarks), and unique_genes (baseline) (**Fig. 2c-e**), performance increases as cells gain a higher proportion of foreign transcripts, with full doublets showing the highest AUROC. This trend is consistent across methods (**Fig. S2**), but the performance decreases systematically as perturbed cells lose a higher proportion of their own transcripts (**Fig. 2f**). Accordingly, doublet methods can detect partial molecular admixture, though sufficient molecules from both cells are needed, highlighting the importance of transcript loss when assessing contamination.

In single-cell RNA-seq, the doublet rate depends on the number of cells loaded, with more cells increasing the probability of co-encapsulation. The doublet methods accept this prior probability as a parameter, including Scrublet and DoubletFinder used for spatial transcriptomic quality control. However, the prior probability of segmentation errors in spatial data likely differs from the doublet base rate. To test whether this prior probability impacts performance, we repeated our earlier experiments – which perturbed 10% of cells – by perturbing 1% and 20% of cells (**Fig. S3-4**), finding that the methods show similar performance across perturbation rates (**Fig. 2g**).

We next evaluated how gene panels impact performance. To this end, we created full doublets using real and simulated panels. cxds2 and scDblFinder.score, for example, show higher performance for actual panels even when the simulated panels are much larger (**Fig. 2h**), such that real panels with ∼500 genes perform better than simulated panels of size 10,000. These patterns are consistent across doublet methods (**Fig. S1d**), suggesting that the current gene panels capture a large portion of the variability (low dimensionality) present in transcriptome-wide expression. Finally, we examined how performance varies between real gene panels given differences in their sizes (98 to 1122) and overlaps (5 to 477). Performance on detecting full doublets is highly consistent (**Fig. 2i**), with the largest standard deviation (0.04) being fairly modest, obviating the need to select specific gene sets for doublet detection.

Our results suggest that doublet methods successfully detect partial molecular admixture, are unaffected by the error base rate, and can effectively leverage extant gene panels, making them promising candidates for quality control in spatial transcriptomics.

### Doublet methods mostly detect partial molecular admixture in spatial transcriptomic data

We assembled 16 spatial transcriptomic studies representing 7 technologies (BARseq, CosMx, EEL FISH, MERFISH/MERSCOPE, Molecular Cartography, STARmap, Xenium), 4 staining strategies (nuclear, cytoplasmic, membranous, or none), and 4 segmentation algorithms (CellPose, ClusterMap, Resolve, Xenium Analyzer) (**Fig. 3a**). The slices analyzed contain 21,002 to 207,591 cells, and all except one are large (full or half) coronal or sagittal sections (**Fig. S5**). The studies show different transcript detection sensitivities based on the technology used. For example, the average expression of shared genes is highly correlated when both studies use the Xenium platform (**Fig. 3b**, Pearson’s R = 0.94), but the correlation is lower when, say, one study uses MERFISH and the other uses EEL FISH (**Fig. 3c**, Pearson’s R = 0.33). In general, higher correlations are seen when studies use the same technology (**Fig. 3d**, Mann-Whitney U Test, Similar ≥ Dissimilar, p-value = 0.035), with low correlations observed only between different technologies. Focusing on MERFISH/MERSCOPE, we see higher correlations when the same staining technique is used (**Fig. 3e**, Mann-Whitney U Test, Similar ≥ Dissimilar, p-value = 0.0086), suggesting that data acquisition and processing parameters can introduce further variability.

**Figure 3.**
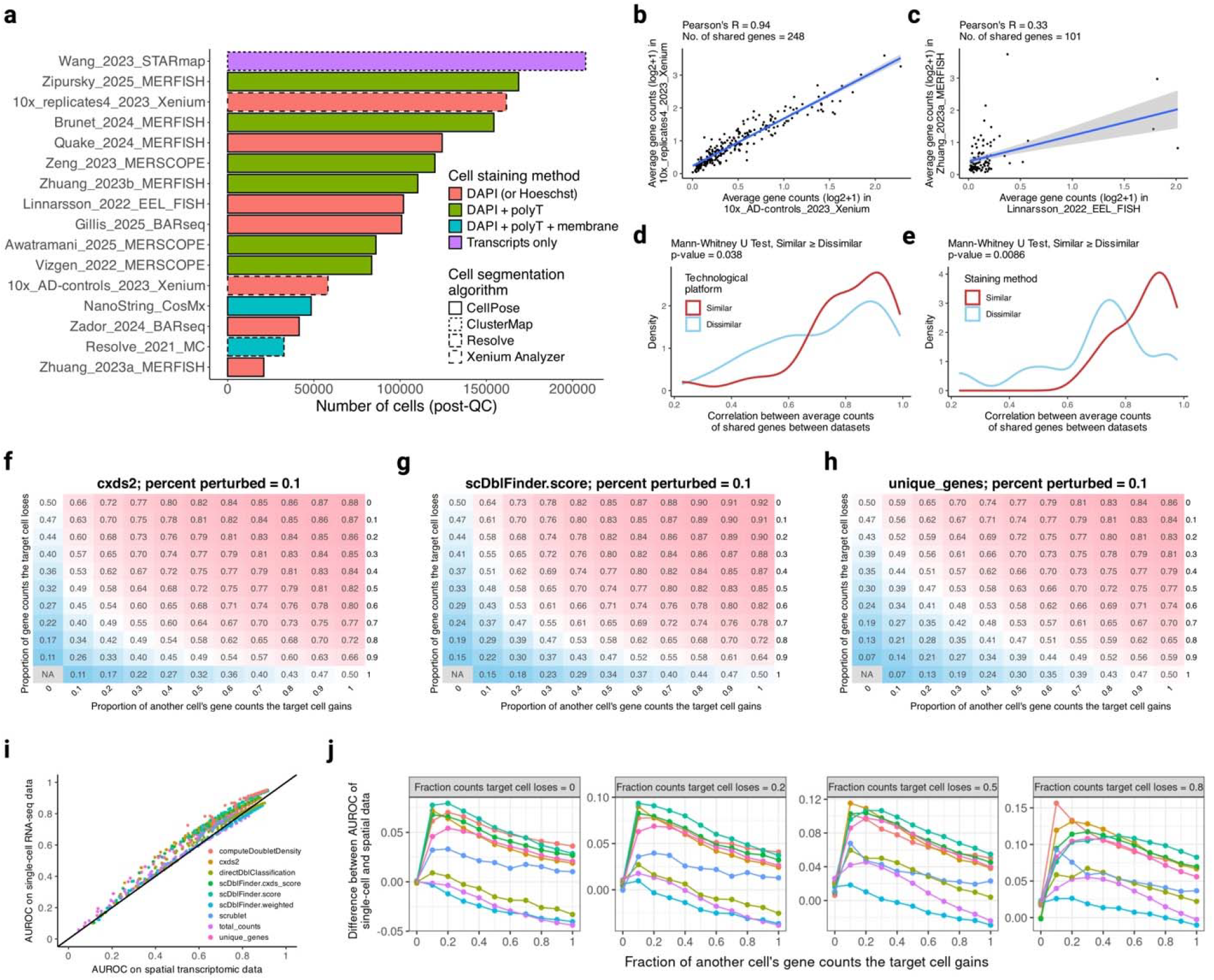
Error detection in spatial transcriptomic data using randomly chosen donor cells. **a**, Datasets (slices) ordered by the number of cells. Colors indicate staining techniques and outlines represent segmentation algorithms. **b**, Transcript detection sensitivity between datasets using the same technology (Xenium). Points are genes’ average counts, so only those genes contained in both panels are included. Blue line and shaded region indicate the regression line and 95% confidence interval, respectively. **c**, Same as in (**b**) but between datasets using different technologies (MERFISH and EEL FISH). **d**, Pearson’s correlations calculated as in (**b**) or (**c**) colored by whether the datasets use the same or different technologies. **e**, Pearson’s correlations as before but between MERFISH/MERSCOPE datasets colored by whether they use the same or different staining methods. Panels (**d**) and (**e**) involve one-sided Mann-Whitney U Test, similar ≥ dissimilar, with p-values 0.038 and 0.0086, respectively. **f-h**, Heatmaps show AUROCs across the perturbation space for cxds2 (**f**), scDblFinder.score (**g**), and unique_genes (**h**). Perturbed cells gain transcripts from randomly chosen donor cells, not from spatial neighbors. Each entry shows AUROC averaged over 16 gene panels. In each case, 10% of cells were perturbed. **i**, AUROC in scRNA-seq against spatial transcriptomic data. Points are corresponding entries in the perturbation space and are colored by methods. Black solid line is y = x. **j**, Delta AUROC (scRNA-seq versus spatial) against admixture severity, faceted by transcript loss. Legend is the same as in (**i**). See also Figures S5-9. Created in Biorender.com.

Systematic differences between spatial platforms require doublet methods to generalize across datasets. Using random donors, we perturbed 10% of cells across the perturbation space and computed AUROCs using methods’ doublet scores. We chose random cells (not neighbors) as donors to match our earlier experimental setup with scRNA-seq. This allowed us to quantify how technical differences impact AUROC, providing an upper-bound on performance in spatial data. Like with scRNA-seq, we see strong performance by cxds2, scDblFinder.score, and unique_genes (**Fig. 3f-h**), with AUROC increasing as cells gain a higher proportion of foreign transcripts. All methods show a similar trend (**Fig. S6**). Consistent with our previous results, perturbing 1% or 20% of cells has minimal impact (**Fig. S7-9**).

Nonetheless, performance is systematically lower in spatial than in scRNA-seq data (**Fig. 3i**). Specifically, it differs more when the errors are subtler. For example, scDblFinder.score has AUROC of 0.8 in scRNA-seq and 0.73 in spatial data when cells gain 20% of donor transcripts. Yet, these scores are 0.94 and 0.92 respectively when detecting full doublets. A similar pattern is observed for most methods (**Fig. 3j**, first panel), showing that milder molecular admixture is harder to detect in spatial data. Additionally, transcript loss decreases performance more in spatial compared to scRNA-seq data. For instance, when a full donor cell is gained but 80% of the perturbed cell is lost, scDblFinder.score’s AUROC is 0.81 in scRNA-seq and 0.72 in spatial data, compared to 0.94 and 0.92 respectively for full doublets. This pattern is seen across methods (**Fig. 3j**, last panel), highlighting that doublet methods are particularly sensitive to transcript loss in spatial transcriptomic data.

Our results indicate that doublet methods detect partial molecular admixture in spatial data, showing robustness to myriad technical differences. Compared to scRNA-seq, the lower performance here might be due to global batch effects and extant segmentation errors, giving an upper-bound on error detection in spatial transcriptomics.

### Error-detection between spatial neighbors depends on transcriptional differences

Cell segmentation errors can lead to transcript misassignment between nearby cells. We therefore modelled such errors as follows: perturbed cells gain their nearest neighbors’ transcripts and lose their own transcripts. After perturbing 10% of cells, we used doublet methods’ scores to calculate AUROCs across the perturbation space and averaged the results over 16 spatial studies. We observe similar general patterns as before: all methods’ performance improves with increasing molecular admixture but declines with transcript loss (**Fig. S10**). Perturbing 1% or 20% of cells has minimal effect (**Fig. S11-13**). cxds2, scDblFinder.score, and unique_genes perform the best, with AUROCs of 0.79-0.81 for full doublets (**Fig. 4a-c**). Importantly, Scrublet – the method most commonly used in spatial data – achieved an AUROC of only 0.66, suggesting that current quality control practices may leave substantial segmentation errors undetected.

**Figure 4.**
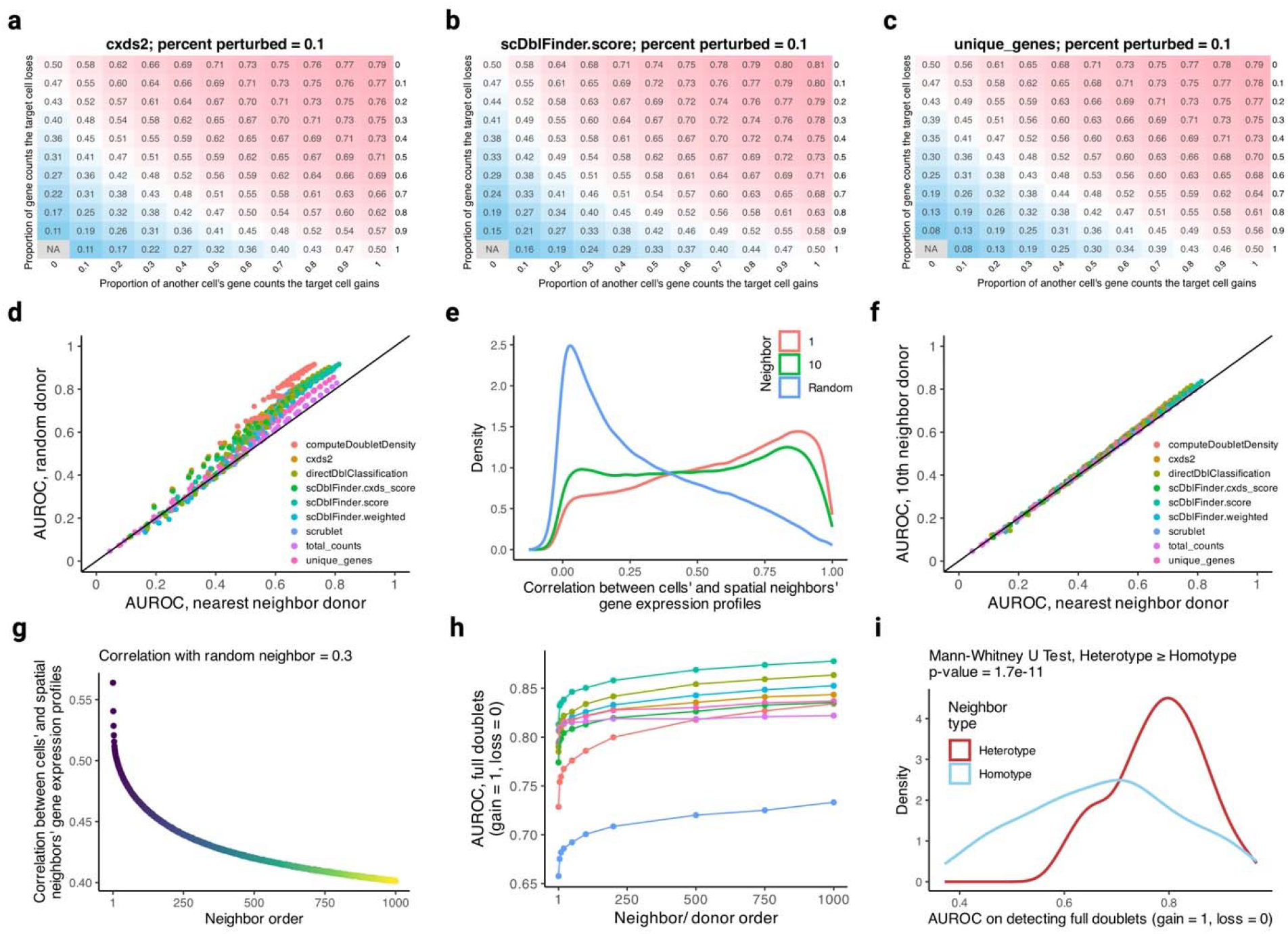
Error detection in spatial transcriptomic data using nearest neighbor donor cells. **a-c**, Heatmaps show AUROCs across the perturbation space for cxds2 (**a**), scDblFinder.score (**b**), and unique_genes (**c**). Perturbed cells gain transcripts from their nearest neighbors. Each entry shows AUROC averaged over 16 gene panels. In each case, 10% of cells were perturbed. **d**, AUROC in spatial data using random versus nearest neighbor donor cells. Points are corresponding entries in the perturbation space and are colored by methods. Black solid line is y = x. **e**, Pearson’s correlations using gene expression profiles of cells and their spatial neighbors (1^st^ or 10^th^) or random cells. Equal number of cell-cell correlations were sampled from each dataset. **f**, Same as in (**d**) but between 10^th^ neighbor and nearest neighbor donor cells. **g**, Pearson’s correlations using gene expression profiles of cells and their spatial neighbors. Each point shows the average calculated using dataset-specific averages. Procedures in (**e**) and (**g**) control for unequal number of cells in different datasets (slices). **h**, AUROC (full doublets) against increasingly distant donor cells. Points represent the sampled donors and are averaged over 16 datasets. Legend is the same as in (**d**) and (**f**). **i**, AUROC (full doublets) colored by whether the nearest neighbor donors are transcriptionally similar or dissimilar to perturbed cells. ‘Homotype’ involves perturbing cells whose cell-neighbor expression correlation (Pearson’s) is in the top 10%. ‘Heterotype’ involves the same relationship when the correlation is in the bottom 10%. Distributions are AUROCs across all methods applied to every dataset. One-sided Mann-Whitney U Test, heterotype ≥ homotype, p-value = 1.7 × 10^−11^. See also Figures S5 and S10-16. Created in Biorender.com.

Molecular admixture between nearby cells, rather than between random ones, more closely approximates cell segmentation errors. We find that the doublet methods’ performance is systematically lower when the donor cells are spatial neighbors (**Fig. 4d**). This may occur due to pre-existing segmentation errors, in which the cells’ observed gene expression profiles are contaminated by their neighbors’ transcripts. This can confound our setup, requiring the doublet methods to discriminate between pre-existing (“normal”) and experimental (“perturbed”) admixtures. To overcome this issue, we chose the 10^th^ neighbor as donor, whose distance from the perturbed cell makes segmentation-led admixture unlikely but preserves the variability in local gene expression (**Fig. 4e**). We repeated our experiments with this change (**Fig. S14-16**), seeing marginal improvement in performance over the nearest neighbor donor (**Fig. 4f**).

However, cells are typically more transcriptionally similar to their nearest neighbor than to their 10^th^ neighbor (see Fig. 4e). We therefore asked whether this difference in expression similarity can explain the observed performance gain. Because transcriptional dissimilarity increases with spatial distance (**Fig. 4g**), we sampled donors at progressively greater distances and found that AUROC increased accordingly (**Fig. 4h**). To distinguish the effects of physical distance from those of expression similarity, we selected nearest neighbor donors conditioned on their gene expression similarity to perturbed cells. Performance was significantly higher for heterotypic than homotypic donors (**Fig. 4i**, Mann-Whitney U Test, Heterotype ≥ Homotype, p-value = 1.7 × 10^-11^), indicating that transcriptional dissimilarity primarily drives performance.

Our results suggest that doublet methods’ performance declines when molecular admixture occurs between spatial neighbors. This happens because nearby cells are transcriptionally similar, making the detection of subtle errors particularly challenging.

### Incorporating single-cell RNA-seq or spatial information does not meaningfully improve performance

Because doublet methods struggle with spatial data, we explored strategies to improve their performance. We first augmented the methods with scRNA-seq data, reasoning that it might allow them to learn the manifold of genuine, segmentation-free cells. To do so, we perturbed spatial cells using their nearest neighbors and concatenated scRNA-seq data in different ratios, inputting the combined expression matrix into doublet methods (**Fig. 5a**). We then used the scores of just the spatial cells to compute AUROCs. Using computeDoubletDensity as an example, we find that the performance is lower when more scRNA-seq cells are included (**Fig. 5b-d**). For instance, the AUROC to detect full doublets is 0.73 when the ratio is 0:1 (spatial-only) or 1:1 but decreases to 0.67 when it is 10:1. This pattern is evident across methods, where performance decreases more with ratio 10:1 than with 1:1 (**Fig. 5e**), indicating that separating perturbed from normal spatial cells becomes *harder* when scRNA-seq cells are included.

**Figure 5.**
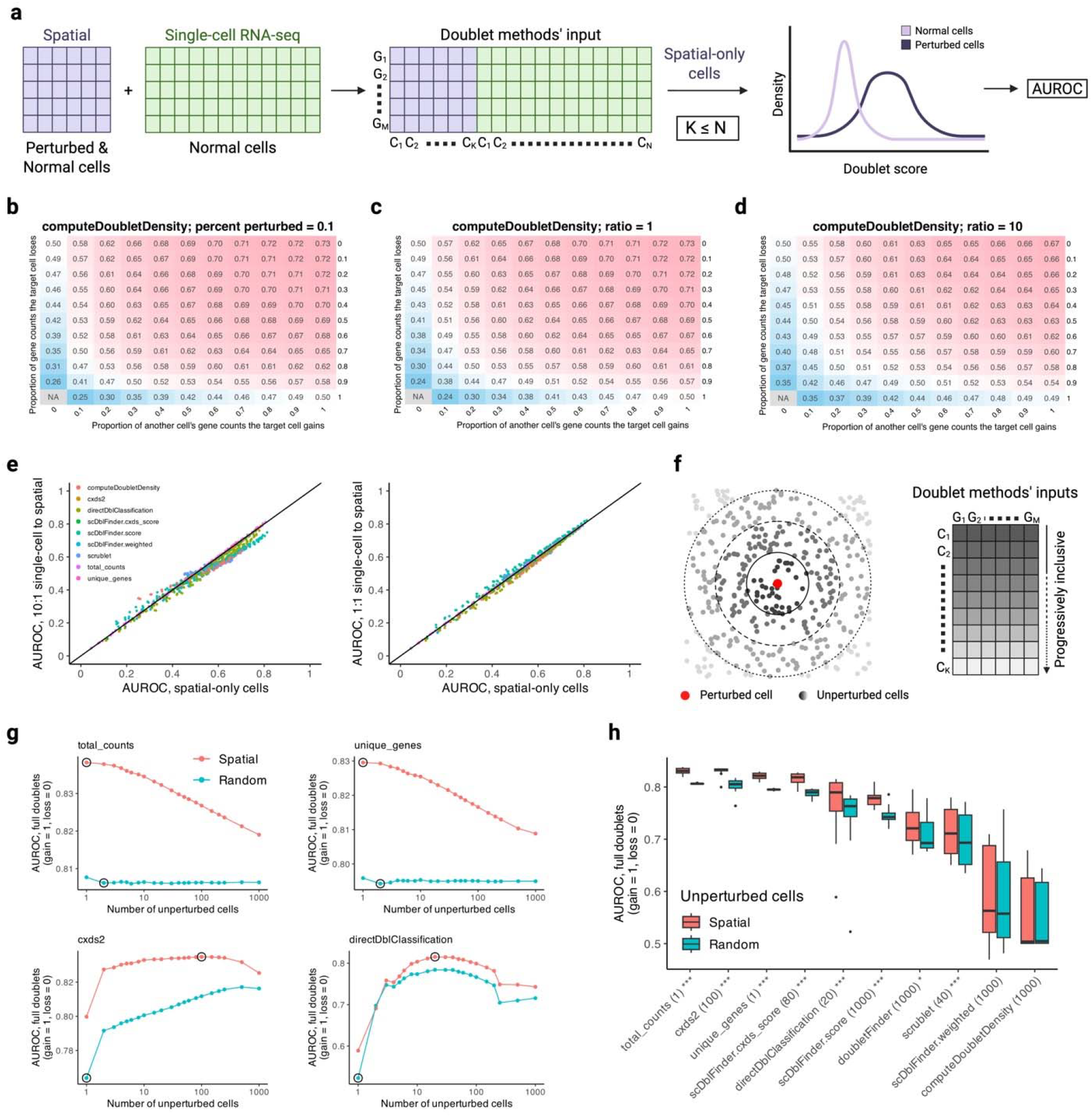
Doublet method augmentation via reference data and spatial locations. **a**, Method augmentation using segmentation-free scRNA-seq data. Left, cells in spatial transcriptomic data are perturbed using their nearest neighbors. The resultant matrix (purple) is combined with normal scRNA-seq data (green) in different ratios. Middle, the combined matrix is inputted into doublet methods. Right, doublet scores of spatial cells are used to calculate AUROC. **b-d**, Heatmaps show computeDoubletDensity’s AUROCs across the perturbation space, with scRNA-seq to spatial ratios of 0:1 (spatial-only) (**b**), 1:1 (**c**), and 10:1 (**d**). Each entry shows AUROC averaged over 16 gene panels. In each case, 10% of cells were perturbed. **e**, Left, AUROC with 10:1 against 0:1 ratio (spatial-only). Right, AUROC with 1:1 against 0:1 ratio. Points are corresponding entries in the perturbation space and are colored by methods. Black solid line is y = x. **f**, Method augmentation using spatial locations. Cells are perturbed using their nearest neighbors and spatial radius is progressively increased to create distinct input matrices. **g**, AUROC (full doublets) against spatial radius, shown for baselines (top) and two methods (bottom). ‘Random’ represents input matrices containing cells from across the tissue. Circles on ‘spatial’ denote maximum performance and on ‘random’ the maximum difference in performance between ‘spatial’ and ‘random’. **h**, Boxplots show AUROC (full doublets) across datasets, grouped by method and colored by whether spatially local or randomly sampled cells are used. Numbers in parentheses (bottom) indicate neighbor order with highest AUROC for ‘spatial’ cells. Boxplots show interquartile range (IQR), center line is median, and whiskers extend to 1.5 × IQR. One-sided paired Wilcoxon signed-rank test, spatial ≥ random, was conducted per method, where * and *** indicate p-values (after Benjamini-Hochberg FDR-correction) less than or equal to 0.05 and 0.001, respectively. See also Figures S5. Created in Biorender.com.

Next, we asked whether spatial information can be used to improve performance. Because nearby cells are transcriptionally similar, we reasoned that restricting the input to these cells may cause the methods to create finer-grained doublet representations. To this end, we perturbed cells using their neighbors and progressively expanded the spatial window (**Fig. 5f**), forcing the methods to sample from a spatially – and transcriptionally – restricted population. Focusing on full doublets, we observe highest AUROCs for total_counts (∼0.84) and unique_genes (0.83) when the window size is 1 (**Fig. 5g**), i.e., input contains the perturbed cell and its second neighbor. (The first neighbor is removed because it is the donor.) The performance here increases because a doublet, which is created by merging a cell with its neighbor, typically contains more total counts and unique genes than the normal cell (2^nd^ neighbor). We also observe improved performance for some doublet methods, e.g., cxds2 and directDblClassification, though the AUROCs are lower than those of our baselines. To assess whether spatially restricted input underlies performance gain, we compared AUROCs when using spatial neighbors versus random cells (**Fig. 5h**, Wilcoxon signed-rank test, Spatial ≥ Random within each neighbor order), finding that most methods perform better when using neighbors.

In short, our augmentation efforts do not meaningfully improve doublet methods’ performance. Nonetheless, their central insights – namely, segmentation-free model of true cells and spatially correlated gene expression – may be leveraged in alternative ways to detect segmentation errors.

### Doublet scores diagnose cell segmentation issues

We have been using doublet scores to identify experimentally perturbed cells. We now ask whether they can diagnose pre-existing cell segmentation issues. To begin, we combined spatial transcriptomic with scRNA-seq data and inputted the matrix into doublet methods. As examples, we use cxds2 (accurate and efficient) and Scrublet (most frequently used in spatial data) scores from Xenium and MERSCOPE, which are well-established and commonly used technologies. Interestingly, in the Xenium data, cxds2 and Scrublet both assign much higher scores to spatial than to scRNA-seq cells (**Fig. 6a**, Spatial > Single-cell, AUROC: cxds2 = 0.84, Scrublet = 0.76), suggesting that spatial data has baseline contamination. By contrast, in the MERSCOPE data, higher scores are assigned to spatial cells by cxds2 but not by Scrublet (**Fig. 6b**, cxds2 AUROC = 0.75, Scrublet AUROC = 0.56). Since cxds2 performed better in our benchmarks, this suggests that Scrublet may overlook segmentation errors when used on spatial data.

**Figure 6.**
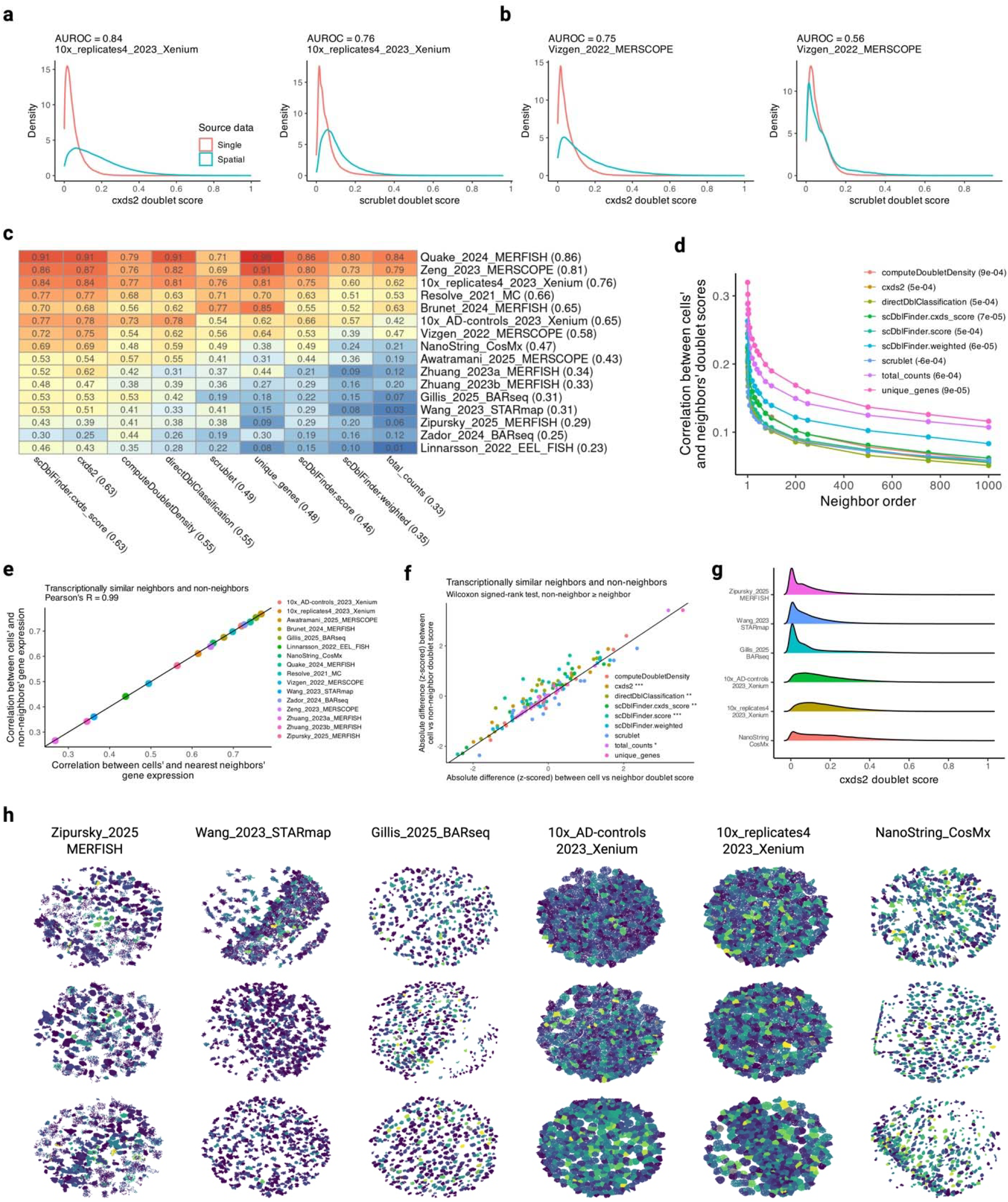
Doublet scores to detect latent cell segmentation issues. **a**, Doublet scores of combined scRNA-seq and spatial transcriptomic data without any perturbations. Colors indicate source datasets, which supply cells in equal amounts. Higher scores predict ‘spatial’ as source data, so AUROC indicates possible contamination in spatial relative to scRNA-seq data. cxds2 (left) and Scrublet (right) scores are shown on a Xenium dataset. **b**, Same as in (**a**) but for a MERSCOPE dataset. **c**, AUROC (as in **a**) of methods against datasets, with numbers in parentheses indicating column-wise and row-wise averages, respectively. **d**, Spearman’s correlation between cells’ and neighbors’ doublet scores against neighbor order. Scores were obtained using only spatial data. Each point is the average of dataset-specific averages. Correlations between random cells’ doublet scores are shown in the legend. **e**, Pearson’s correlation using gene expression of cells and non-neighbors against cells and nearest neighbors. Non-neighbors are physically distant cells with high transcriptional similarity to nearest neighbors. Each point represents a dataset-specific average. **f**, Delta of cell versus non-neighbor doublet scores against cell versus nearest neighbor doublet scores. Each point is the average of dataset-specific averages and is colored by doublet method. One-sided paired Wilcoxon signed-rank test, non-neighbor ≥ neighbor, was conducted per method, where (next to method names) *, **, and *** indicate p-values (after Benjamini-Hochberg FDR-correction) less than or equal to 0.05, 0.01, and 0.001, respectively. Black solid lines in (**e**) and (**f**) are y = x. **g**, cxds2 doublet scores for datasets containing transcript-level information. Transcript coordinates, gene identity, and cell assignment yielded gene expression matrices, which were input into cxds2. **h**, Example transcripts shown in space and colored by cxds2 scores (from **g**) of cells that contain them. Columns (datasets) are ordered by doublet scores and rows (fields-of-view, FOV) by transcript count. Every FOV has 500 cells. Color scale is shared between and is comparable across plots. See also Figures S6. Created in Biorender.com.

Next, we asked which spatial datasets show elevated doublet scores. Using our earlier scheme, we computed AUROCs, where higher scores predict “spatial” (rather than “scRNA-seq”) as the source dataset. Focusing on cxds2, we find that Quake, Zeng, and the two Xenium datasets show high AUROCs (**Fig. 6c**), indicating potentially higher baseline errors. Others like Zador and Zipursky show low AUROCs, which occur when spatial cells have lower doublet scores, possibly revealing conservative segmentation strategies. Many datasets have AUROCs around 0.5, indicating globally reliable cell segmentation. These observations are consistent across methods. Crucially, our inferences here depend on treating the reference scRNA-seq as ground truth: deviations from this null indicate possible contamination or overly stringent segmentation.

We now assess doublet scores at the level of individual cells. Here, we use spatial data as input to doublet methods, finding that nearby cells have more similar scores than distant cells (**Fig. 6d**, Nearest neighbor vs Random cells, Spearman’s rho = ∼0.3 vs. ∼0). Since gene expression is spatially correlated, we asked whether doublet scores remain spatially coupled once transcriptional similarity is controlled. To answer this question, we identified distant cells (non-neighbors) with very similar gene expression profiles to nearest neighbors, such that the cell-neighbor and cell-non-neighbor relations were virtually identical (**Fig. 6e**, Pearson’s R = 0.99), thus perturbing spatial relations while maintaining expression similarity. We find that for some methods, like cxds2 and scDblFinder.score, neighbors have more similar doublet scores (**Fig. 6f**), highlighting that they can identify spatially structured contamination, e.g., due to cell segmentation, even when gene expression – their sole input – is explicitly controlled.

Lastly, we spatially visualize the cells using their doublet scores. Six of our datasets had transcript-level information, including spatial coordinates, gene identity, and cell assignment (post-segmentation). This allowed us to create gene expression matrices, which we used to compute cxds2 doublet scores. Consistent with our previous results, some datasets have higher doublet scores with right-skewed distributions (**Fig. 6g**). Using spatial coordinates and cell assignments, we plotted the transcripts and colored them by their cells’ doublet scores (**Fig. 6h**), such that spatial contiguity and score intensity visually reveal distinct cells. Interestingly, the datasets with high doublet scores, such as Xenium and CosMx, have cells that appear densely packed together and contain little space between them, possibly leading to transcript misassignment at cell boundaries.

Our results indicate that high doublet scores likely reflect spatially correlated baseline contamination. Importantly, spatial datasets with high scores exhibit visual properties associated with segmentation issues, making doublet methods appropriate diagnostic tools in spatial transcriptomics.

## Discussion

A major goal in genomics is to understand the molecular composition of cells. A natural approach is therefore to capture intact cells and sequence their mRNA transcripts, as in scRNA-seq^73,74^. Imaging-based spatial transcriptomics reverses this logic: transcripts are first detected in situ, after which they are assigned to cells through segmentation^2^. The definition of a “cell” – and which molecules it contains – therefore becomes computational rather than purely biological. Doublet methods have consequently been used to identify potentially problematic cells because they detect expression profiles containing transcripts from more than one cell^31–36^. Their use in spatial data implicitly assumes that molecular admixture arising from scRNA-seq doublets resembles that produced by segmentation errors. Although plausible, this assumption has not been systematically tested, leaving unclear whether methods designed to detect complete admixture can also identify the subtler, partial admixture produced by segmentation errors^27–30^.

This study modeled cell segmentation errors as partial molecular admixture and evaluated doublet methods under a variety of conditions. Methods performed well across gene panels, error frequencies, detection chemistries, staining strategies, and segmentation algorithms. cxds2 and scDblFinder.score reliably detected strong admixture between nearby cells, while the number of unique genes provided a similarly strong simple baseline. Importantly, all methods performed better when adjacent cells were transcriptionally distinct. This is encouraging because admixture between different cell types is particularly likely to introduce misleading expression profiles, consistent with the emphasis of doublet methods on detecting heterotypic doublets in scRNA-seq^31–36^. Doublet scores also revealed underlying segmentation issues, including dense transcript point clouds and minimal separation between adjacent cells. We observed this, for example, in Xenium data, whose segmentation strategy prioritizes transcript retention^12^.

Beyond identifying problematic cells, our results suggest several ways in which doublet scores could improve spatial transcriptomic data quality^27–30,37–41,63–72^. Algorithms that support fine-tuning from user annotations could use high-scoring cells to prioritize difficult regions for manual segmentation. This could form an iterative procedure in which cells are segmented, high-scoring regions are manually corrected, and the algorithm is subsequently retrained or re-applied. Doublet scores could similarly complement transcript reassignment methods that redistribute molecules between neighboring cells to reduce admixture. Their spatial structure may help identify local regions requiring reassignment and provide an orthogonal measure of post-reassignment quality. Finally, because some segmentation errors will inevitably remain, doublet scores provide a principled basis for filtering admixed cells rather than relying solely on count-based thresholds, which may remove large but transcriptionally pure cells while retaining smaller admixed cells.

Several limitations should be considered. Our perturbation framework models segmentation errors as transcript loss and admixture from neighboring cells. Although this captures a central consequence of incorrect cell boundaries, real segmentation errors may be more complex, involving multiple neighboring cells, irregular boundaries, or other forms of transcript misassignment. Moreover, because doublet methods operate on gene expression profiles, segmentation errors that produce little molecular admixture may remain difficult to detect. Finally, although we evaluated diverse spatial technologies and datasets^44–59^, our analysis focused primarily on mouse brain tissue, and performance may differ in tissues with distinct cellular organization or transcriptional diversity.

Overall, our results support the use of doublet methods for detecting cell segmentation errors in spatial transcriptomics. We particularly recommend cxds2^33^ because it performed well while scaling readily to large datasets of approximately 200,000 cells and 1,000 genes. An alternative is scDblFinder.score^36^ if computational resources allow. More broadly, our results support partial molecular admixture as a useful framework for modeling segmentation errors and show that methods originally developed for scRNA-seq doublets can provide a scalable approach for identifying such errors in spatial transcriptomic data.

## Resource availability

### Lead contact

Requests for further information and resources should be directed to and will be fulfilled by the lead contact, Jesse Gillis.

### Materials availability

This study did not generate new reagents.

### Data and code availability

- This paper analyzes existing, publicly available data, which are accessible via the associated publications or websites^44–59^.
- All original code has been deposited at Zenodo and is available at https://doi.org/10.5281/zenodo.22744799^75^ as of the date of publication.
- Any additional information required to reanalyze the data reported in this paper is available from the lead contact upon request.

## Acknowledgements

A.S. acknowledges funding support from NSERC Canada Graduate Scholarship, Ontario Graduate Scholarship (OGS), and University of Toronto FAST Doctoral Fellowship. J.G. acknowledges funding support from NSERC Discovery RGPIN-2025-06198.

## Author contributions

Conceptualization – AS and JG. Formal analysis – AS and JG. Methodology – AS. Software – AS. Funding acquisition – JG. Supervision – JG. Writing, original draft – AS. Writing, review and editing – AS and JG.

## Methods

### Perturbation strategy

We model cell segmentation errors as partial molecular admixture, where the perturbed cell loses its transcripts and gains another cell’s transcripts. To do so, we apply binomial thinning to cell counts. Let **x**= (*x*_l_,*x*_2_,…,*x*_*g*_) be a cell with *g* genes, where *x*_*j*_ ∈ {0,1,2,…}. Binomially thinning x with probability *p* treats the counts of each gene independently. For gene *j, x*′_*j*_ |*x*_*j*_ ∼ Binomial(*x*_*j*_ ,*p*) yields the thinned cell **x**′ = (*x*′_1_ ,*x*′_2_ ,…,*x*′_*g*_). For example, if *x*_*j*_ = 12 and *p*= 0.5, each of the 12 transcripts is independently treated with probability 0.5, namely *x*′_*j*_ ∼ Binomial(12,0.5), generating *x*′_*j*_ ∈ {0,1,2,…,12} rather than exactly 12 × 0.5 = 6. That is, each molecule is independently chosen with *p*= 0.5, so between 0 to 12 gene *j* molecules may be selected. Importantly, *p* represents the expected fraction of each gene‘s molecules selected, since *E*[*x*′_*j*_ |*x*_*j*_ ] = *px*_*j*_ , and thus it also represents the expected fraction of the cell’s total molecules selected, since if 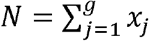, then 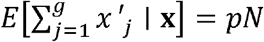. This strategy therefore has the same average effect as naïve scaling *p***x**, but it preserves non-negative integer counts, avoids arbitrary rounding, and introduces sampling variability expected when transcripts are independently selected, e.g., lost or gained during segmentation.

We use binomial thinning to create gain-loss perturbations. Let 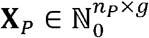 denote the expression matrix of cells randomly selected for perturbation and 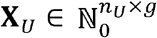 for the unperturbed cells. In practice, we set 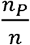 as 0.01, 0.1, or 0.2. For each perturbed cell *i*,, let *d*(*i*) denote its paired donor in 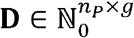, with **D**_*i*_ =**x**_*d*(*i*)_. Transcript loss and gain are defined as L|**X**_*P*_ ∼ Binomial(**X**_*P*_,*p*_loss_) and **G**|**D** ∼ Binomial(**D**,*p*_gain_). We independently vary *p*_loss_ and *p*_gain_ from 0 to 1 in increments of 0.1 and, once determined, keep their values fixed within an experiment. The perturbed cells are thus 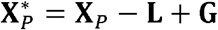, and the final expression matrix is 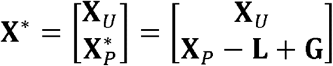. In our implementation, **X**^*^ contains unperturbed and perturbed cells in random order rather than as stacked, as shown here. A binary indicatory vector **z** ∈{0,1}^*n*×l^ tracks the cells’ perturbation status.

### Donor cells and structured noise

Perturbed cells receive transcripts from donor cells. This can involve molecular transfer, where donors effectively lose their transcripts, or duplication, where transcripts are added to perturbed cells without affecting donors. Both cases introduce structured artifacts: molecular transfer directly perturbs donor profiles, whereas duplication artificially increases the similarity between perturbed cells and their donors. We therefore exclude donor cells from the final matrix **X**^*^.

Our approach to donor exclusion depends on whether they are random cells or spatial neighbors. For random donors, we construct a separate donor pool such that each perturbed cell is paired with exactly one donor, and each donor is used at most once. For spatial donors, the donating neighbor is removed from **X**^*^ so that it does not remain as an unperturbed (“normal”) cell. Importantly, in cell-to-neighbor mapping, the same cell may be the nearest neighbor of two or more cells. In such cases, the donating neighbor is allowed to perturb just one of these cells.

### Doublet detection methods

We ran all doublet detection methods using default parameters unless stated otherwise. The input in all cases was **X**^*^.

We ran the following methods using scDblFinder^36,76^ (v1.22.0): computeDoubletDensity, cxds2, directDblClassification, scDblFinder.cxds_score, scDblFinder.score, and scDblFinder.weighted. cxds2 was originally published as part of scds^33,77^, and its implementation in scDblFinder differs only slightly from the original. Although related, cxds2 and scDblFinder.cxds_score are distinct methods.

We used DoubletCollection^35,78^ (v1.1.0) to run Scrublet^32,79^, allowing us to execute its python scripts in R. The default number of principal components (PCs) was 30. For datasets containing fewer genes, we reduced this to the largest number of PCs for which Scrublet could be run: 19 for Gillis_2025_BARseq^57^ (133 genes) and Quake_2024_MERFISH^54^ (157 genes) and 14 for Resolve_2021_Molecular_Cartography^45^ (99 genes) and Zador_2024_BARseq^55^ (109 genes). For the experiment reported in Figure 5h, we used 10 PCs across all datasets.

We ran DoubletFinder^31,80^ (v2.0.3) following its standard Seurat^81,82^ (v5.4.0) pre-processing workflow. Counts were processed using NormalizeData, variable features were identified using the vst method, and expression values were scaled using ScaleData before principal component analysis (PCA). All genes were retained for PCA, as the requested number of variable features exceeded the number of genes assayed in our spatial datasets. DoubletFinder was run using PCs 1-10, *p*_*N*_ = 0.25, and a fixed *p*_*K*_ = 0.09 across datasets. We modified the DoubletFinder code to return continuous pANN scores underlying its binary doublet classifications, enabling calculation of AUROCs (see below). Due to its memory requirements, DoubletFinder could not be run on several large spatial datasets. Its performance is therefore not reported for experiments in which these requirements were prohibitive.

All methods assign a doublet score to each cell. We used these scores together with the perturbation labels to calculate AUROCs^83–86^, with higher doublet scores taken to predict perturbed cells.

### Data preprocessing

We obtained scRNA-seq data from Allen Institute’s whole-brain mouse atlas^60^, in which individual files correspond to distinct brain regions. We pooled cells together from across regions and randomly partitioned them into batches of ∼50,000 cells. This approximates the average number of cells in our spatial slices and represents transcriptional heterogeneity across the brain. All experiments were performed on these batches.

We obtained spatial transcriptomic datasets from a variety of sources (see Data and code availability)^44–59^. Some datasets contained a single slice, typically near the center of the brain, whereas others contained multiple slices. For the latter, we selected high-quality central slices that lacked obvious artifacts, such as tissue folds, damage, or air bubbles. The final collection therefore comprises large, heterogeneous tissue sections spanning both cortical and subcortical regions.

Spatial datasets reported genes using either gene symbols or Ensembl IDs, e.g., *Gad1* or ENSMUSG00000070880. To standardize gene identifiers, we converted gene symbols to Ensembl IDs using mappings provided by the Allen Institute atlas^52^. This facilitated gene matching across spatial datasets and between spatial and scRNA-seq data. Where required, we subset the scRNA-seq data to the corresponding spatial gene panel and ensured identical gene ordering before concatenating the expression matrices. A small number of genes could not be matched in three datasets: NanoString_CosMx^44^ (942 of 950 genes matched), Vizgen_2022_MERSCOPE^46^ (478 of 483), and Resolve_2021_Molecular_Cartography^45^ (98 of 99). The unmatched genes likely reflect minor naming discrepancies in gene symbols. Analyses involving scRNA-seq used the matched gene sets, whereas spatial-only analyses retained the full gene sets. Figures 2a and S1a-b report the smaller, matched gene sets.

Spatial expression matrices also contained negative probes, codewords, and other non-genic features, which were excluded from all analyses. Across all experiments, we removed cells with fewer than 20 total counts or fewer than 5 detected genes.

### Doublet method augmentation using spatial information

We tested whether doublet detection could be improved by restricting the input matrix to the spatial neighborhood of each perturbed cell. Each iteration contained a single perturbed cell, with all other cells remaining unperturbed, and the neighboring donor cell was removed as before. We repeated this procedure until 10% of cells were considered as perturbed and report the mean AUROC across iterations.

### Cell-neighbor correlations

We compute two types of cell-neighbor correlations using either gene expression or doublet scores. Cell-neighbor pairs are identified using RANN’s function nn2^87–89^, which returns the indices of each cell’s neighbor at a specified neighbor rank.

Let **X**∈ ℝ ^*n*x*g*^ denote the cell-by-gene expression matrix and *d*(*i*) the spatial neighbor of cell ,. Using the neighbor indices, we construct the row-aligned neighbor matrix **X**_*N*_ ∈ ℝ ^*n*x*g*^, (**X**_*N*_)_*i*·_ = **X**_*d*_(_*i*_)_·_, such that row *i* of **X**_*N*_ contains the expression profile of cell *i*’s neighbor. Correlation is then calculated only between corresponding rows, *r*_*i*_ = cor(**X**_*i*·_,(**X**_*N*_)_*i*·_)), *i*= 1,…,n, yielding **r**= (*r*_l_,… ,*r*_*n*_) ∈ [-1,1]^*n*^. Thus, correlations are calculated only for cell-neighbor pairs rather than for all pairs of cells.

Gene expression provides a vector of measurements for each cell, allowing a separate correlation to be calculated for every cell-neighbor pair. By contrast, each doublet score is a scalar, so an analogous per-pair correlation cannot be computed. We therefore calculate the correlation across all cell-neighbor score pairs. Let **s**= (*s*_l_,…,*s*_*n*_) ∈ ℝ^*n*^ denote the vector of doublet scores and construct the neighbor-aligned vector *s*_*N*_ = (*s*_*d*(l)_,…,*s*_*d*(*n*)_) ∈ ℝ ^*n*^. The cell-neighbor score association is *r*_*s*_= cor(**s**,**s**_*N*_), giving a single correlation coefficient describing the association between doublet scores of cells and their spatial neighbors.

### Dataset-balanced estimates

Spatial datasets ranged from around 20,000 to over 200,000 cells. Because larger datasets contribute more cell-level observations, we first averaged within each dataset (slice) and then across datasets, thereby giving each dataset equal weight. This prevents larger datasets and their dataset-specific properties from disproportionately influencing the overall estimate.

### Neighbor replacement with non-neighbors

We tested whether nearby cells have similar doublet scores after controlling for transcriptional similarity. For each cell, we identified its nearest spatial neighbor and a matched non-neighbor. Candidate non-neighbors were drawn from the 11th to 1000th nearest cells by spatial distance. From these candidates, we selected the non-neighbor whose expression profile was most similar to that of the nearest neighbor, thereby replacing each neighbor with a transcriptionally matched non-neighbor. We then calculated the absolute difference in doublet scores between the focal cell and its nearest neighbor and, separately, between the focal cell and its matched non-neighbor.

## Supplemental information

Document S1. Supplementary Figures S1-S16.

