## Supplementary Figures for "Detecting cell segmentation errors using doublet methods"

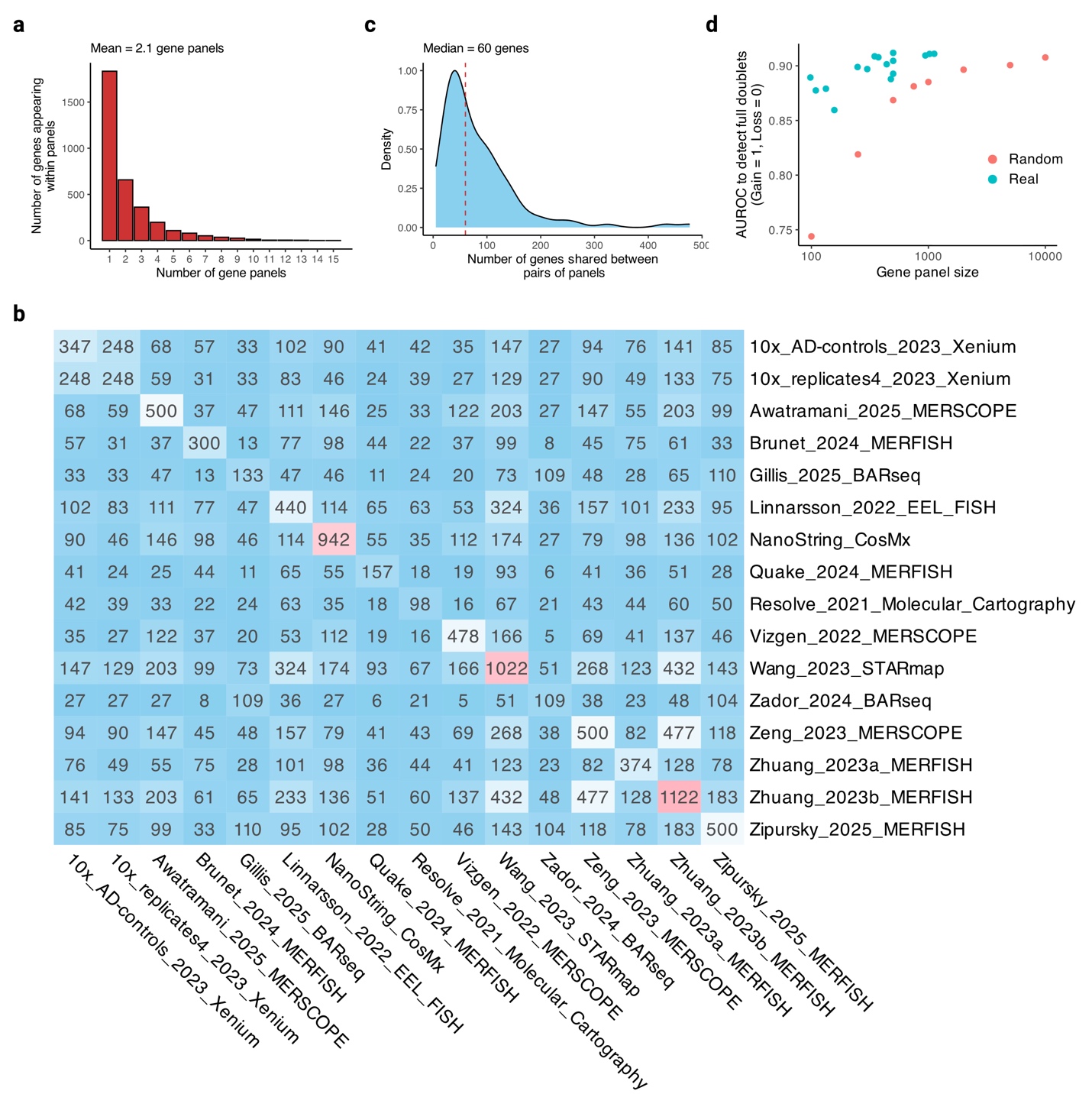

**Figure S1. Gene panel characterization, related to Figure 2. a**, Presence of genes across panels. **b**, Number of genes shared between panels. Diagonal shows panel sizes. See methods for details. **c**, Number of genes shared between panels (from **b**), with red dashed line as median. **d**, AUROC (full doublets) against gene panel size for real and simulated (random) panels, with results averaged across methods. Data points for simulated panels are the average across 100 runs. Created in Biorender.com.

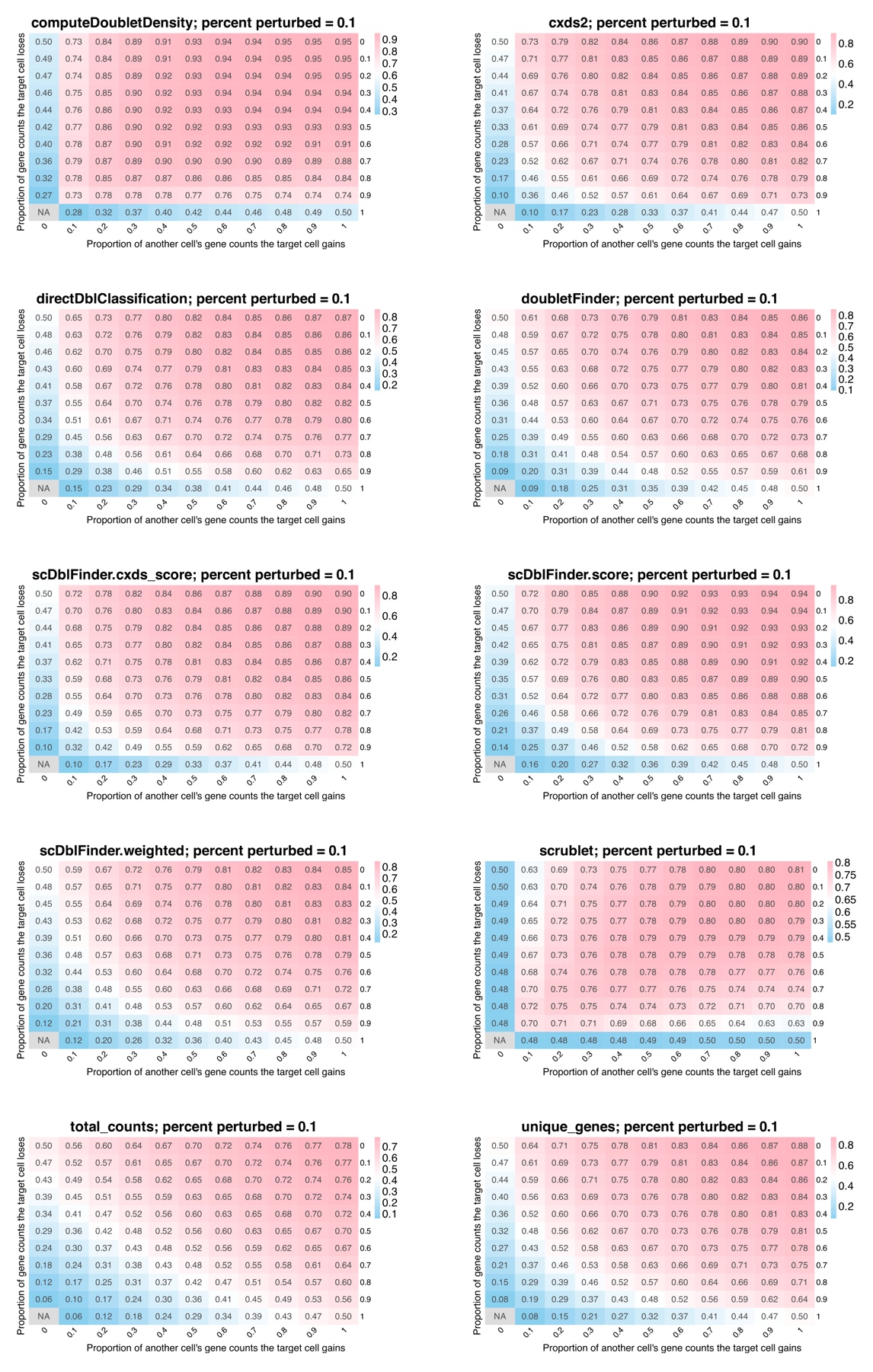

**Figure S2. Error detection in scRNA-seq data, with 10% perturbation rate, related to Figure 2.** Heatmaps show AUROCs across the perturbation space for all methods. Each entry shows AUROC averaged over 16 gene panels. In each case, 10% of cells were perturbed. Created in Biorender.com.

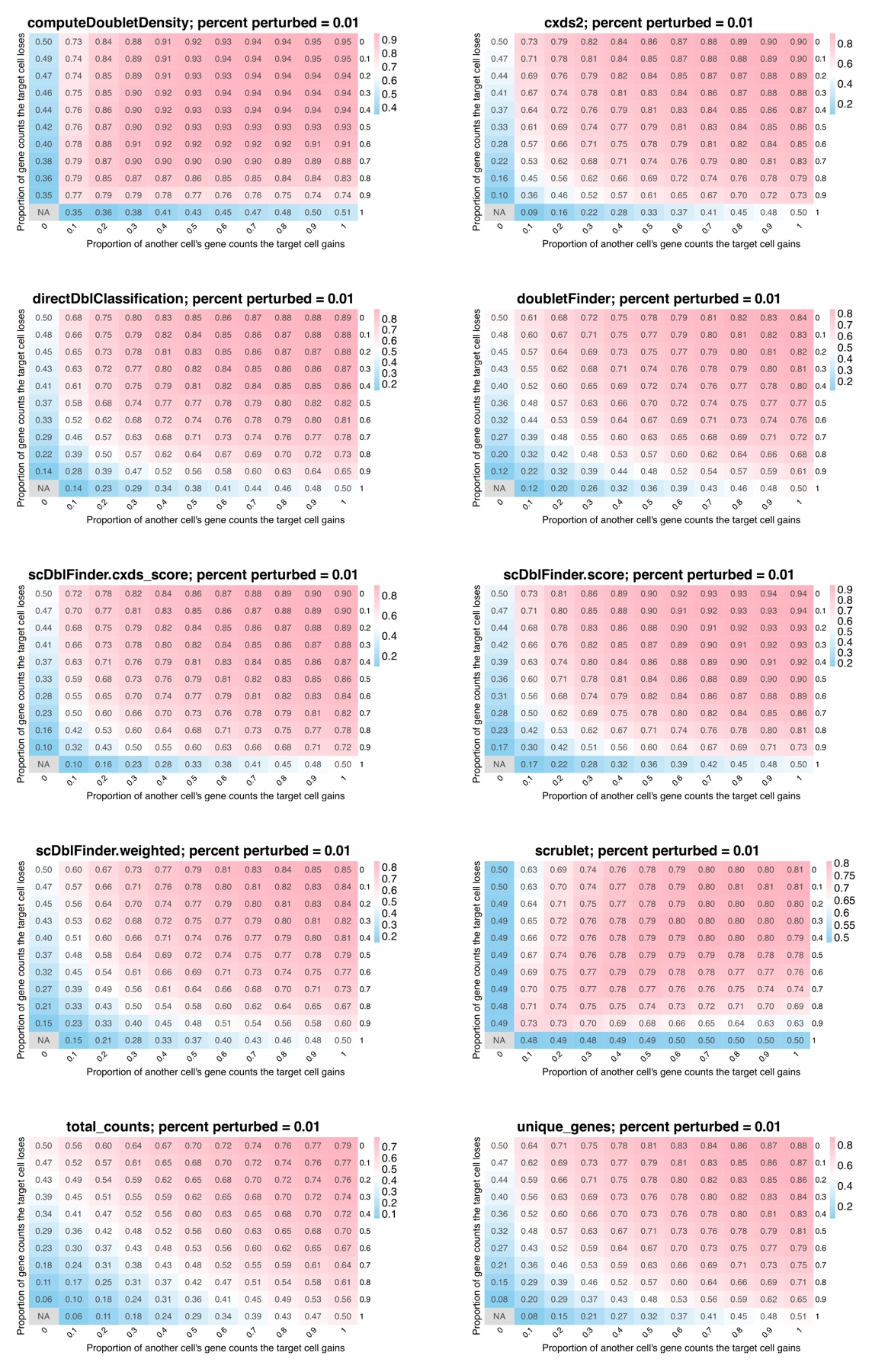

**Figure S3. Error detection in scRNA-seq data, with 1% perturbation rate, related to Figure 2.** Heatmaps show AUROCs across the perturbation space for all methods. Each entry shows AUROC averaged over 16 gene panels. In each case, 1% of cells were perturbed. Created in Biorender.com.

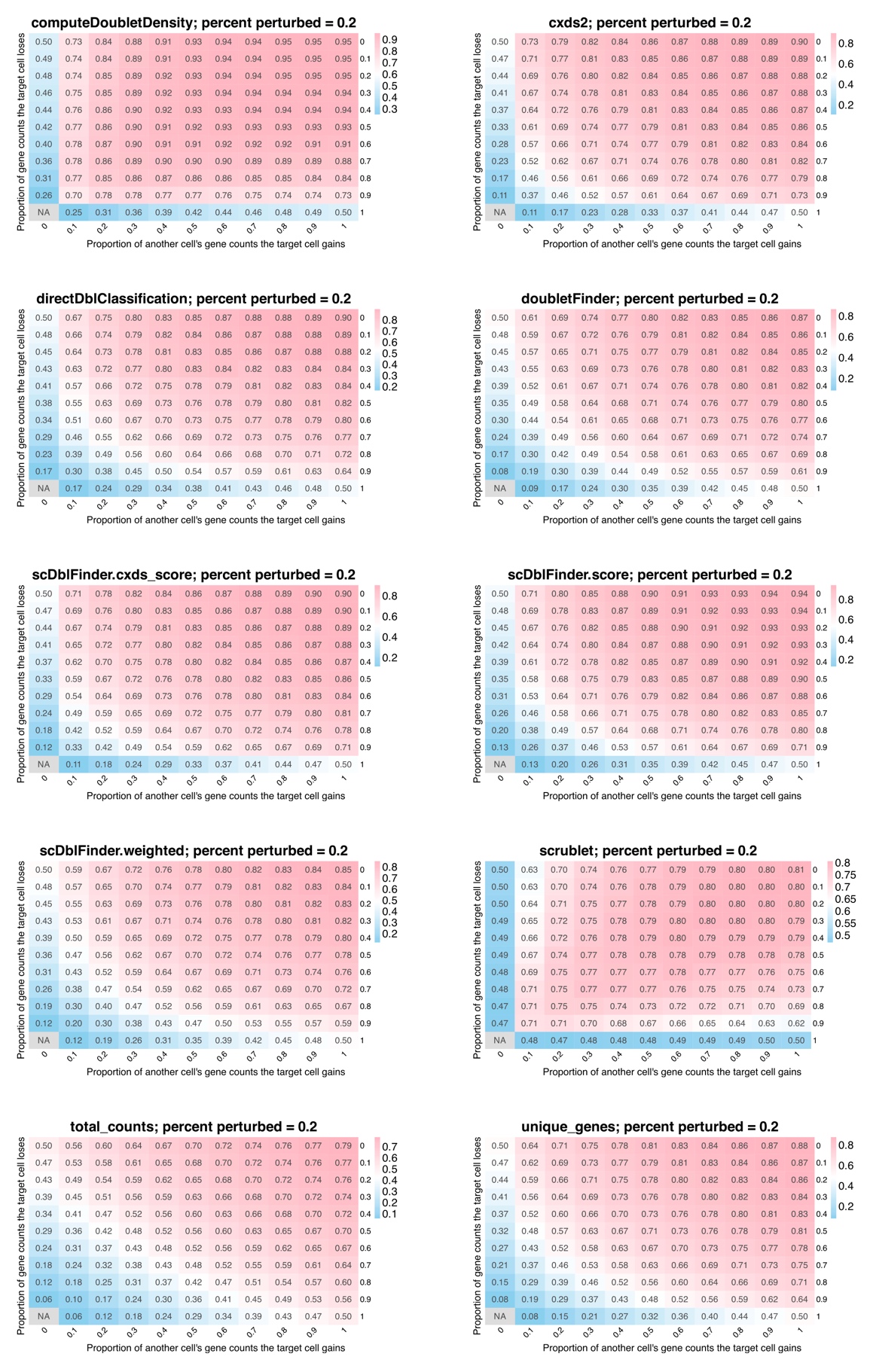

**Figure S4. Error detection in scRNA-seq data, with 20% perturbation rate, related to Figure 2.** Heatmaps show AUROCs across the perturbation space for all methods. Each entry shows AUROC averaged over 16 gene panels. In each case, 20% of cells were perturbed. Created in Biorender.com.

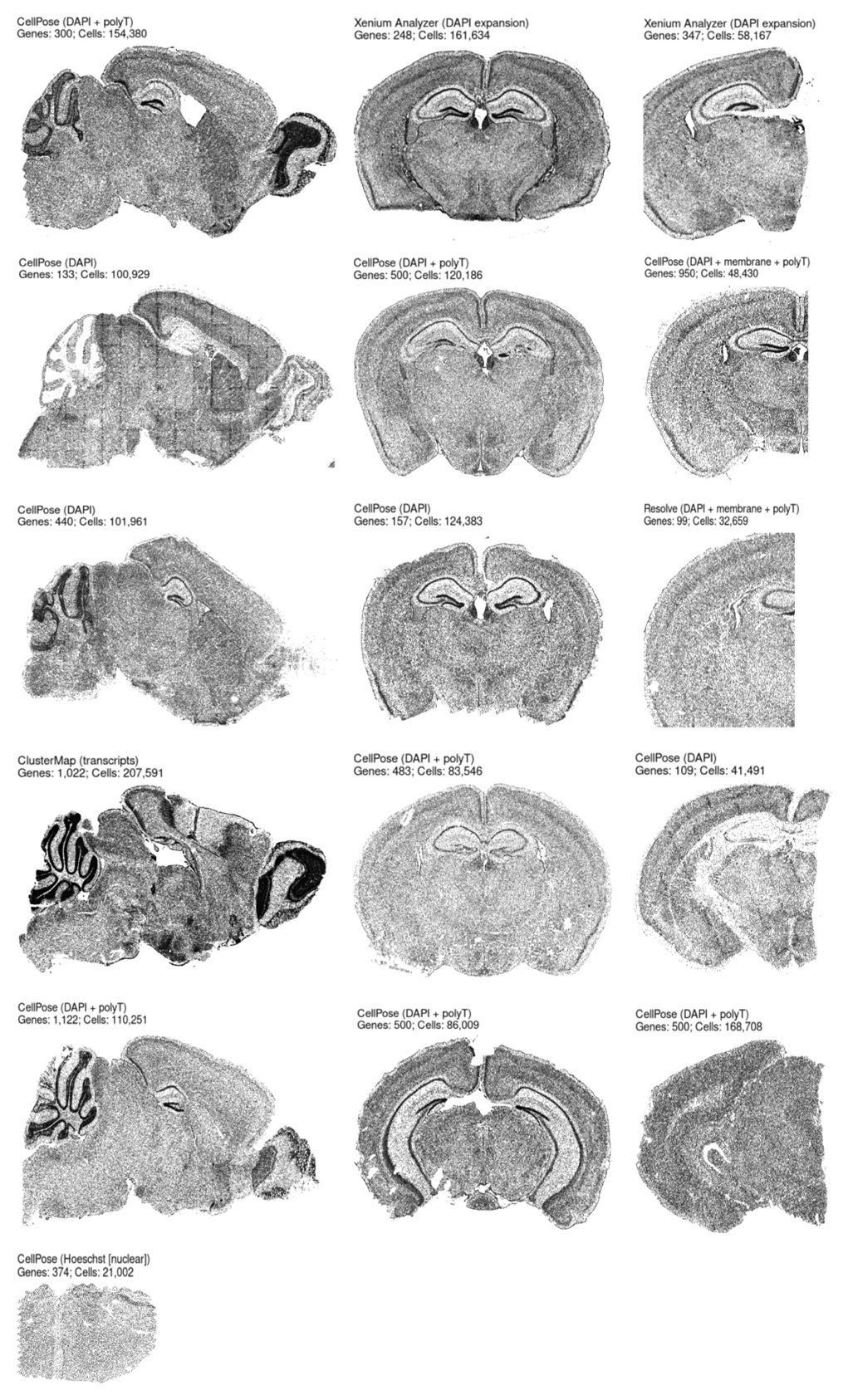

**Figure S5. Spatial transcriptomic slices used, related to Figures 3-6.** High-quality slices chosen from each dataset, with the segmentation algorithm, staining technique, number of genes, and number of cells shown. The datasets are as follows: Brunet_2024_MERFISH, 10x_replicates4_2023_Xenium, 10x_AD-controls_2023_Xenium, Gillis_2025_BARseq, Zeng_2023_MERSCOPE, NanoString_CosMx, Linnarsson_2022_EEL_FISH, Quake_2024_MERFISH, Resolve_2021_Molecular_Cartography, Wang_2023_STARmap, Vizgen_2022_MERSCOPE, Zador_2024_BARseq, Zhuang_2023b_MERFISH, Awatramani_2025_MERSCOPE, Zipursky_2025_MERFISH, and Zhuang_2023a_MERFISH. Created in Biorender.com.

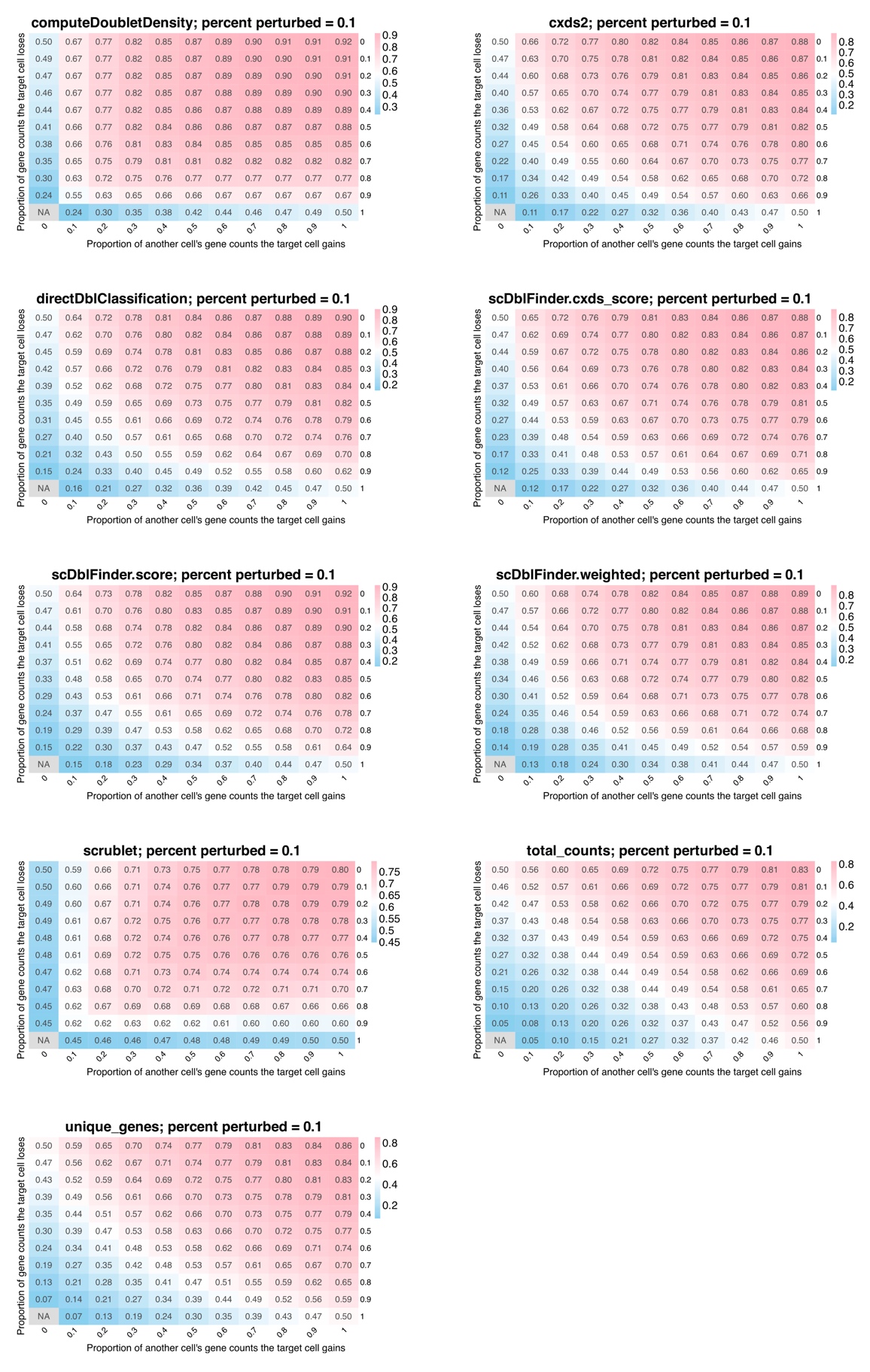

**Figure S6. Error detection in spatial transcriptomic data using random donor cells, with 10% perturbation rate, related to Figure 3.** Heatmaps show AUROCs across the perturbation space for all methods. Each entry shows AUROC averaged over 16 gene panels. In each case, 10% of cells were perturbed. Created in Biorender.com.

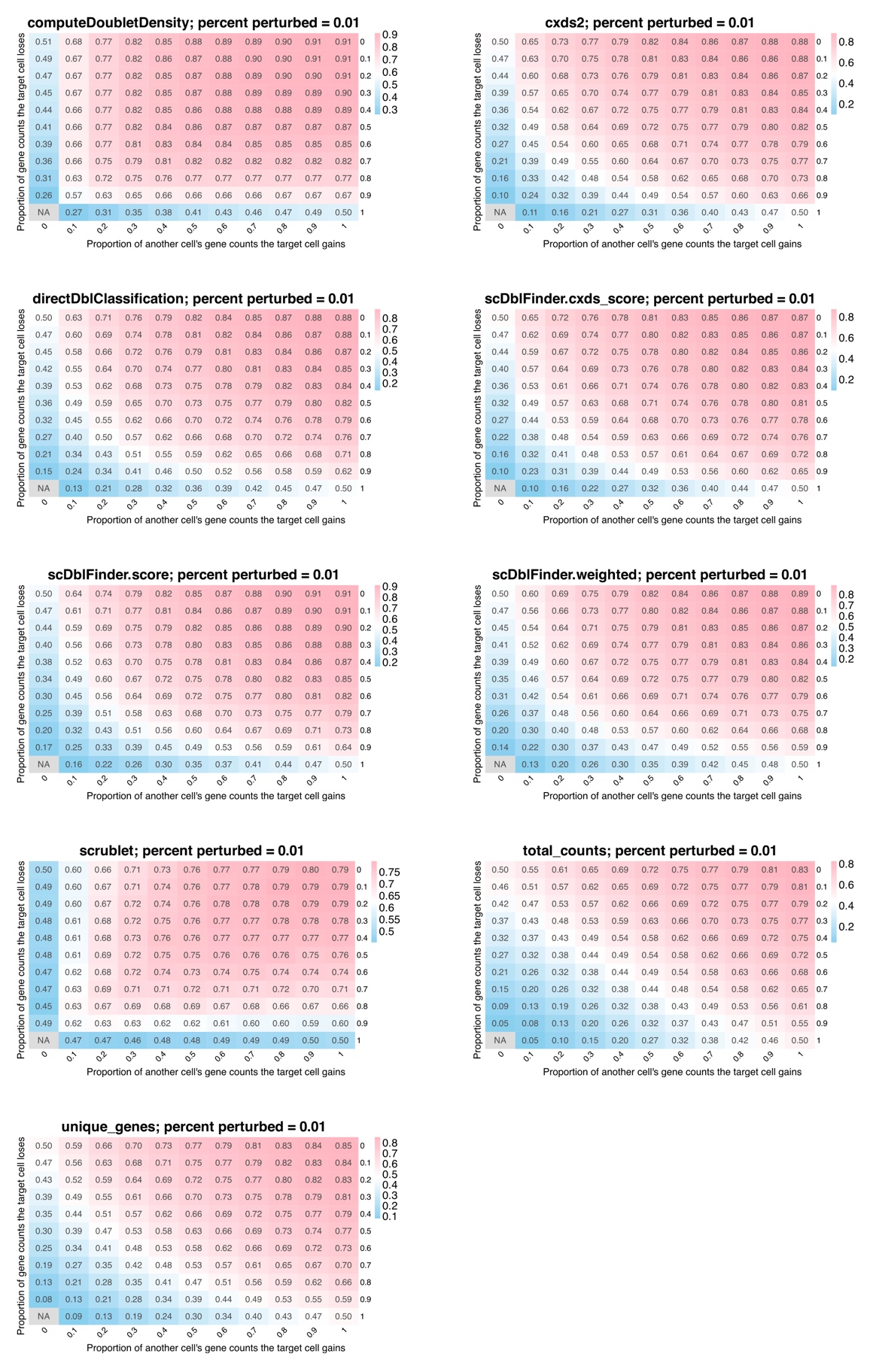

**Figure S7. Error detection in spatial transcriptomic data using random donor cells, with 1% perturbation rate, related to Figure 3.** Heatmaps show AUROCs across the perturbation space for all methods. Each entry shows AUROC averaged over 16 gene panels. In each case, 1% of cells were perturbed. Created in Biorender.com.

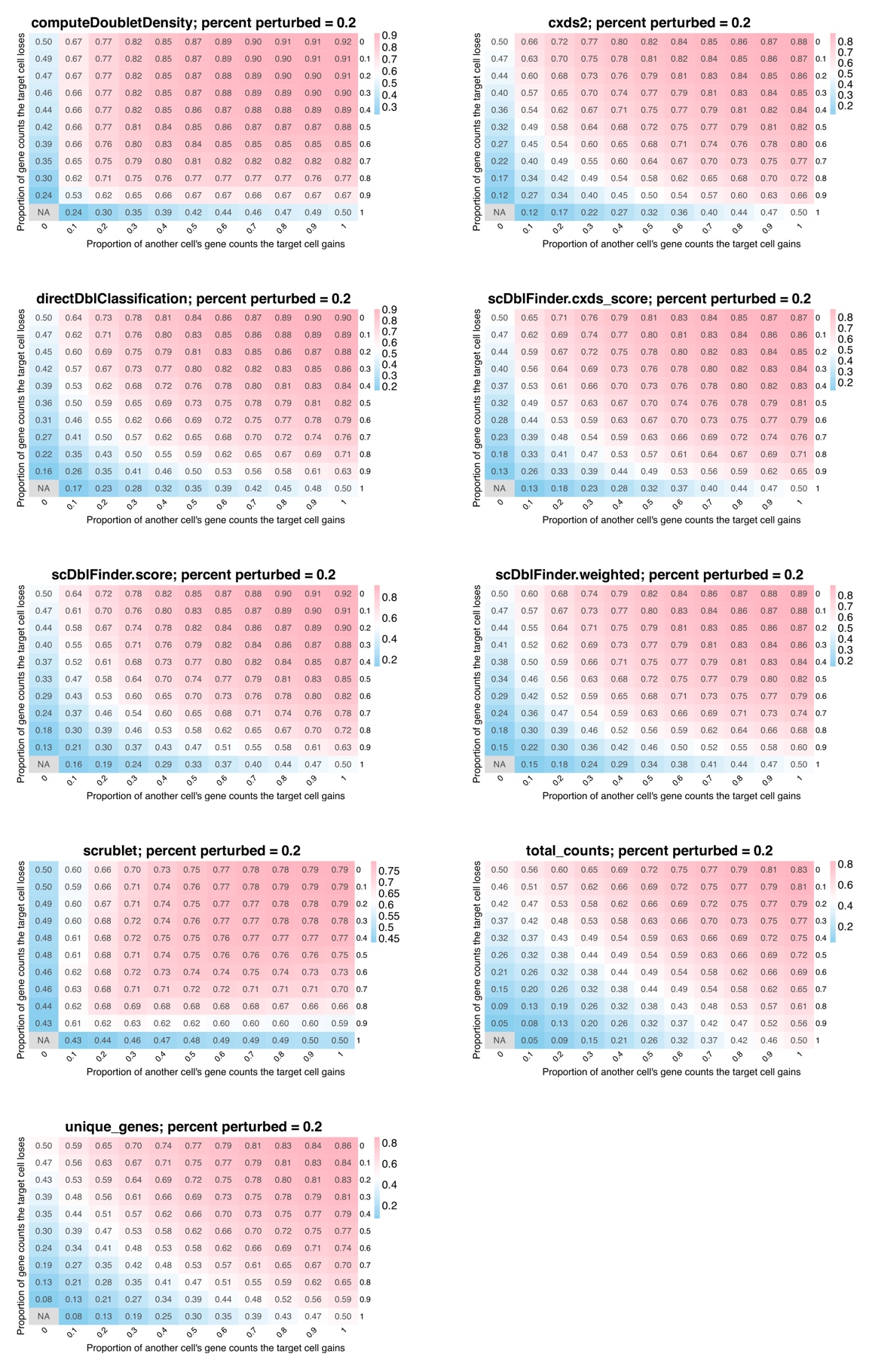

**Figure S8. Error detection in spatial transcriptomic data using random donor cells, with 20% perturbation rate, related to Figure 3.** Heatmaps show AUROCs across the perturbation space for all methods. Each entry shows AUROC averaged over 16 gene panels. In each case, 20% of cells were perturbed. Created in Biorender.com.

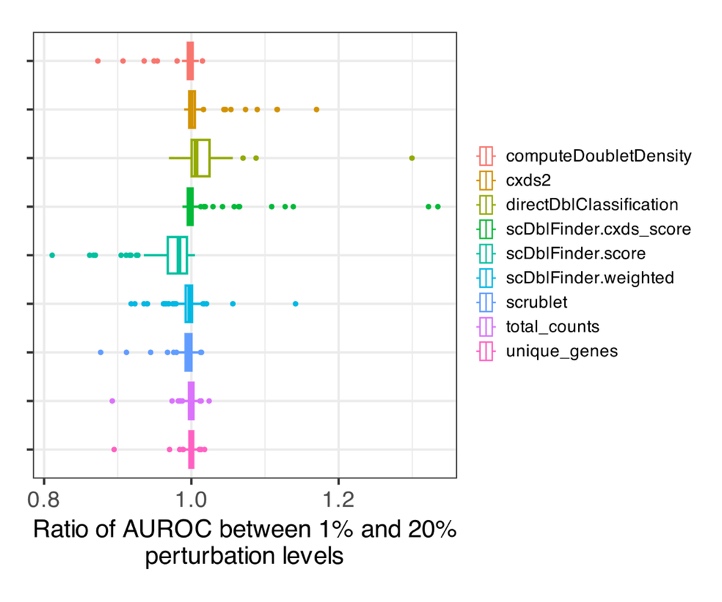

**Figure S9. Perturbation rate comparison in spatial transcriptomic data using random donor cells, related to Figure 3.** Ratio of AUROC when 1% and 20% of cells are perturbed. Ratios are taken between corresponding entries in the perturbation space. See also Figures S7-S8. Created in Biorender.com.

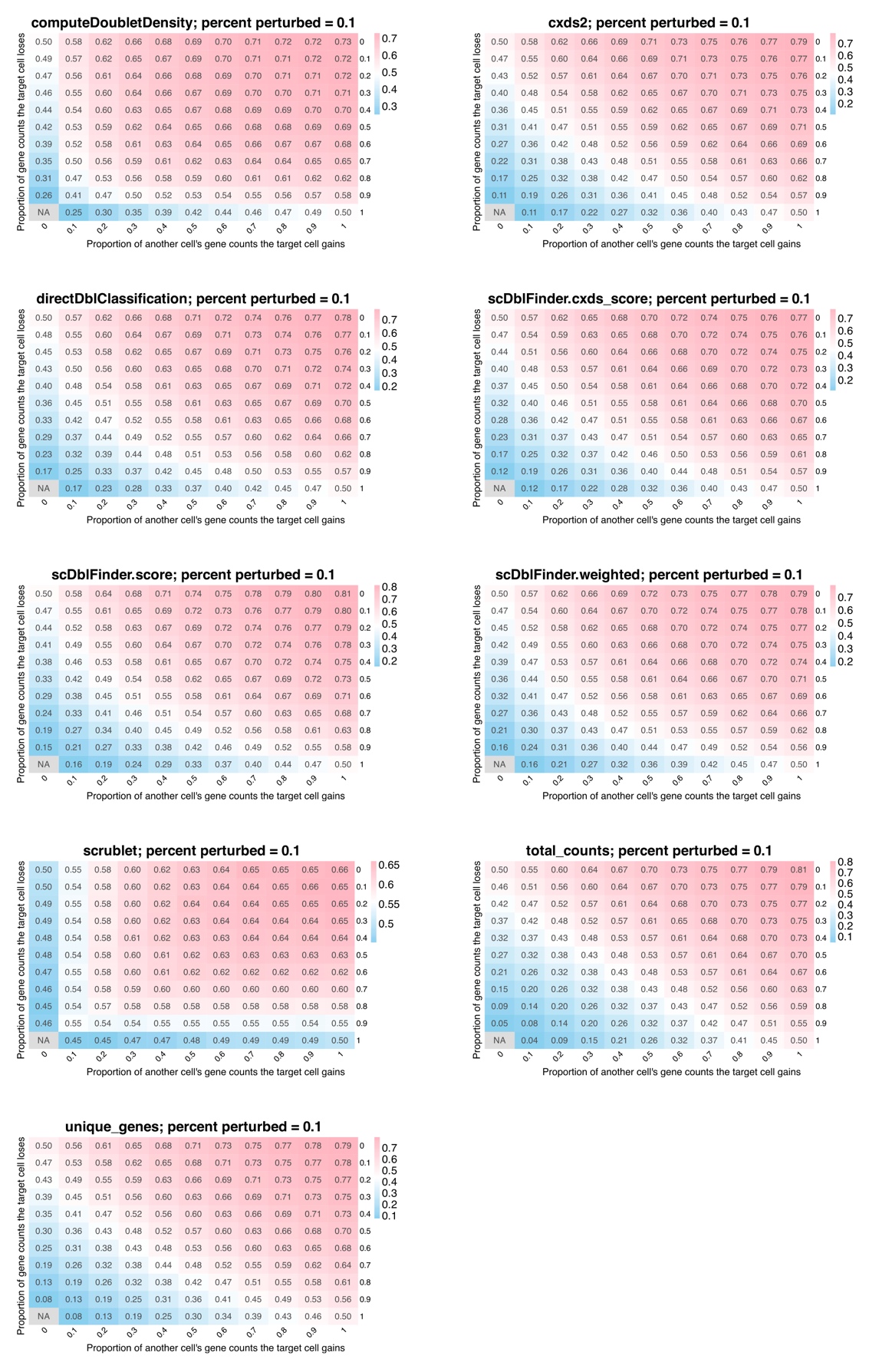

**Figure S10. Error detection in spatial transcriptomic data using nearest neighbor donor cells, with 10% perturbation rate, related to Figure 4.** Heatmaps show AUROCs across the perturbation space for all methods. Each entry shows AUROC averaged over 16 gene panels. In each case, 10% of cells were perturbed. Created in Biorender.com.

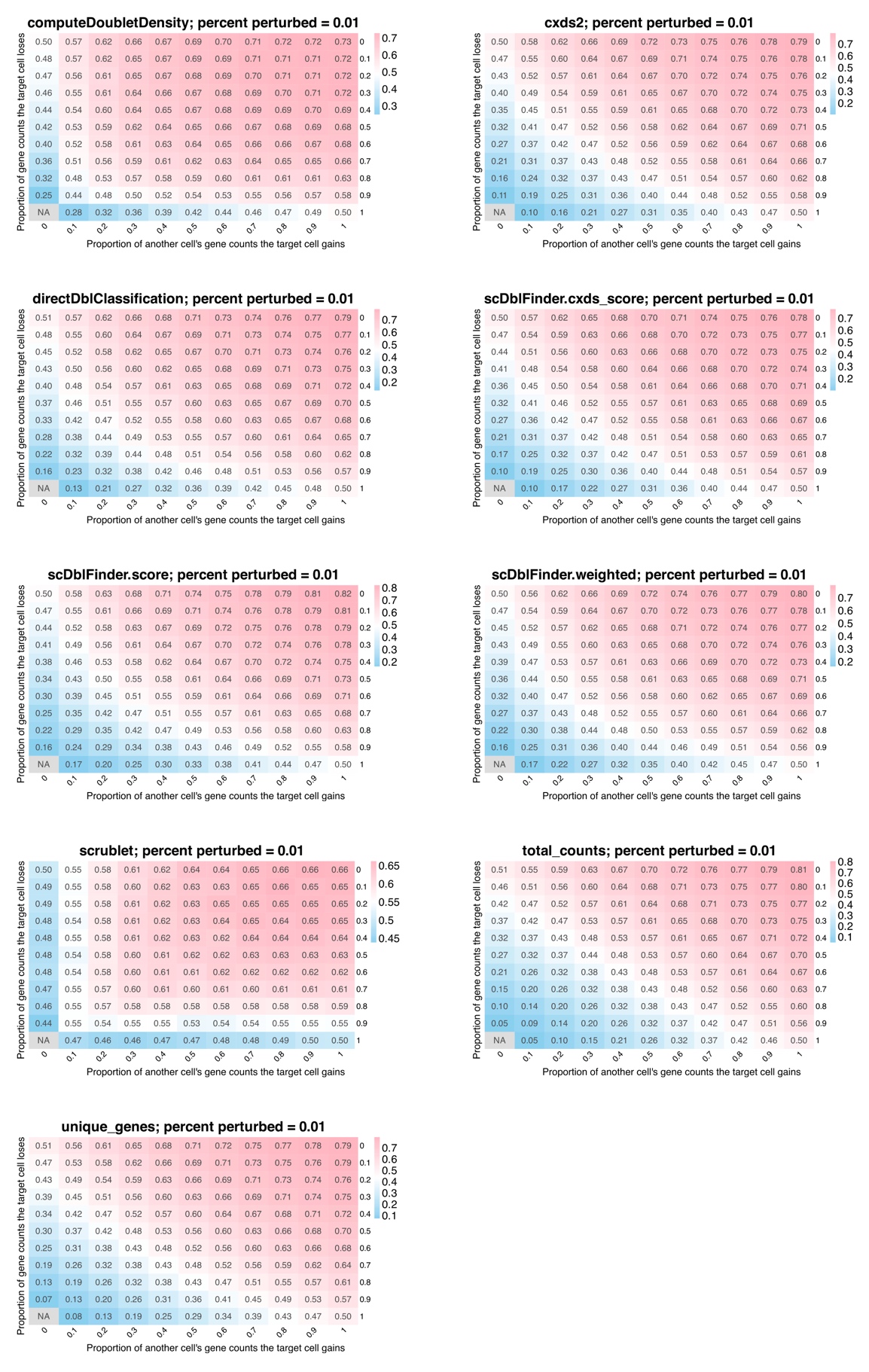

**Figure S11. Error detection in spatial transcriptomic data using nearest neighbor donor cells, with 1% perturbation rate, related to Figure 4.** Heatmaps show AUROCs across the perturbation space for all methods. Each entry shows AUROC averaged over 16 gene panels. In each case, 1% of cells were perturbed. Created in Biorender.com.

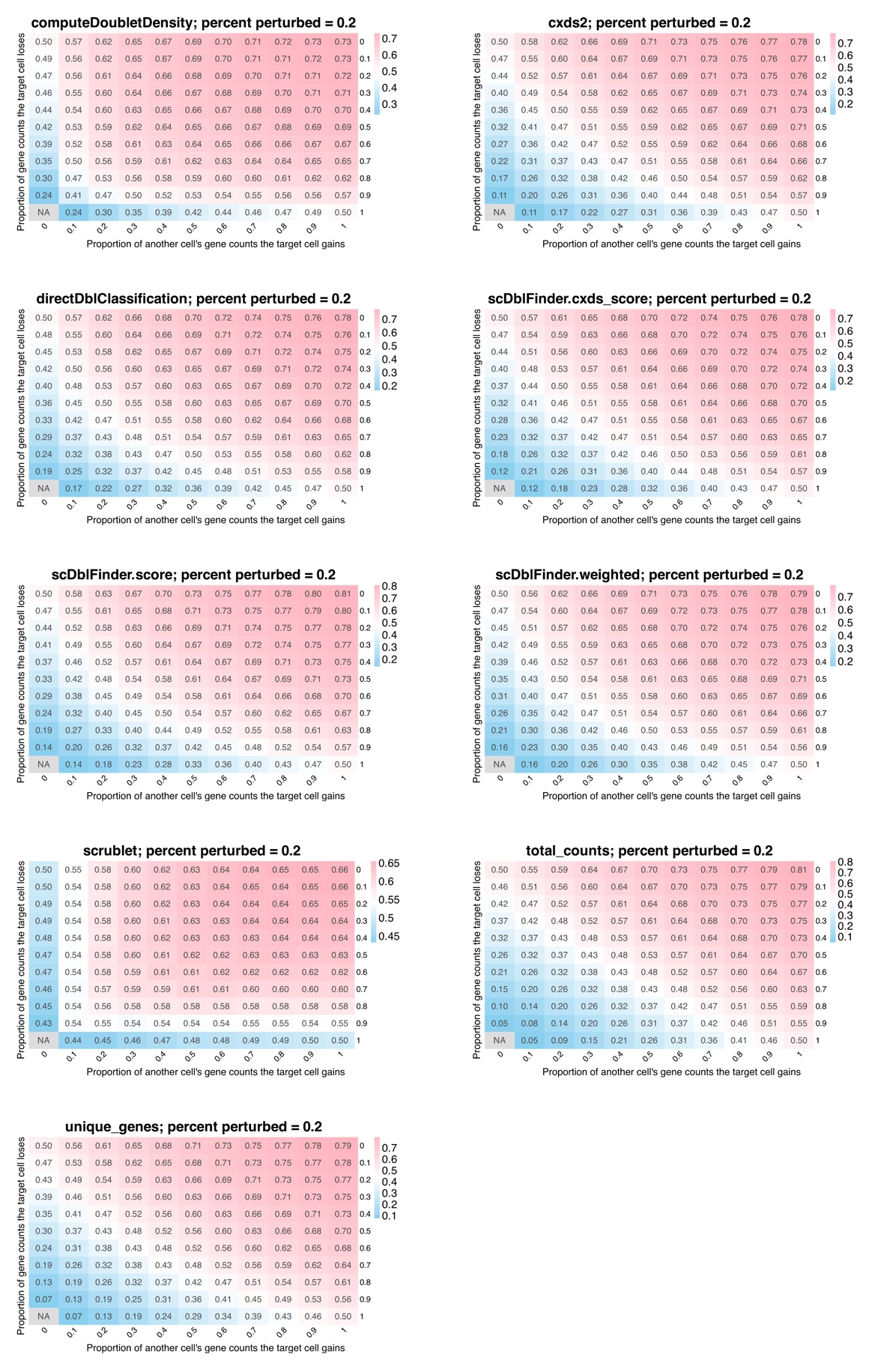

**Figure S12. Error detection in spatial transcriptomic data using nearest neighbor donor cells, with 20% perturbation rate, related to Figure 4.** Heatmaps show AUROCs across the perturbation space for all methods. Each entry shows AUROC averaged over 16 gene panels. In each case, 20% of cells were perturbed. Created in Biorender.com.

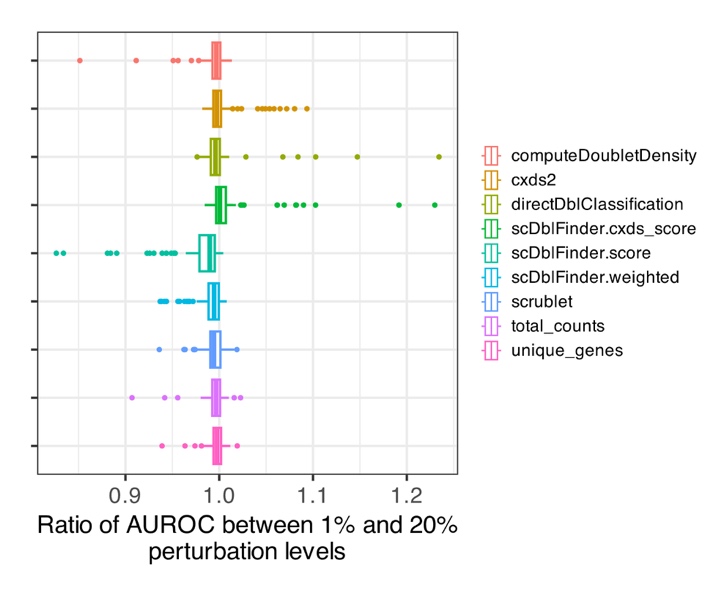

**Figure S13. Perturbation rate comparison in spatial transcriptomic data using nearest neighbor donor cells, related to Figure 4.** Ratio of AUROC when 1% and 20% of cells are perturbed. Ratios are taken between corresponding entries in the perturbation space. See also Figures S11-S12. Created in Biorender.com.

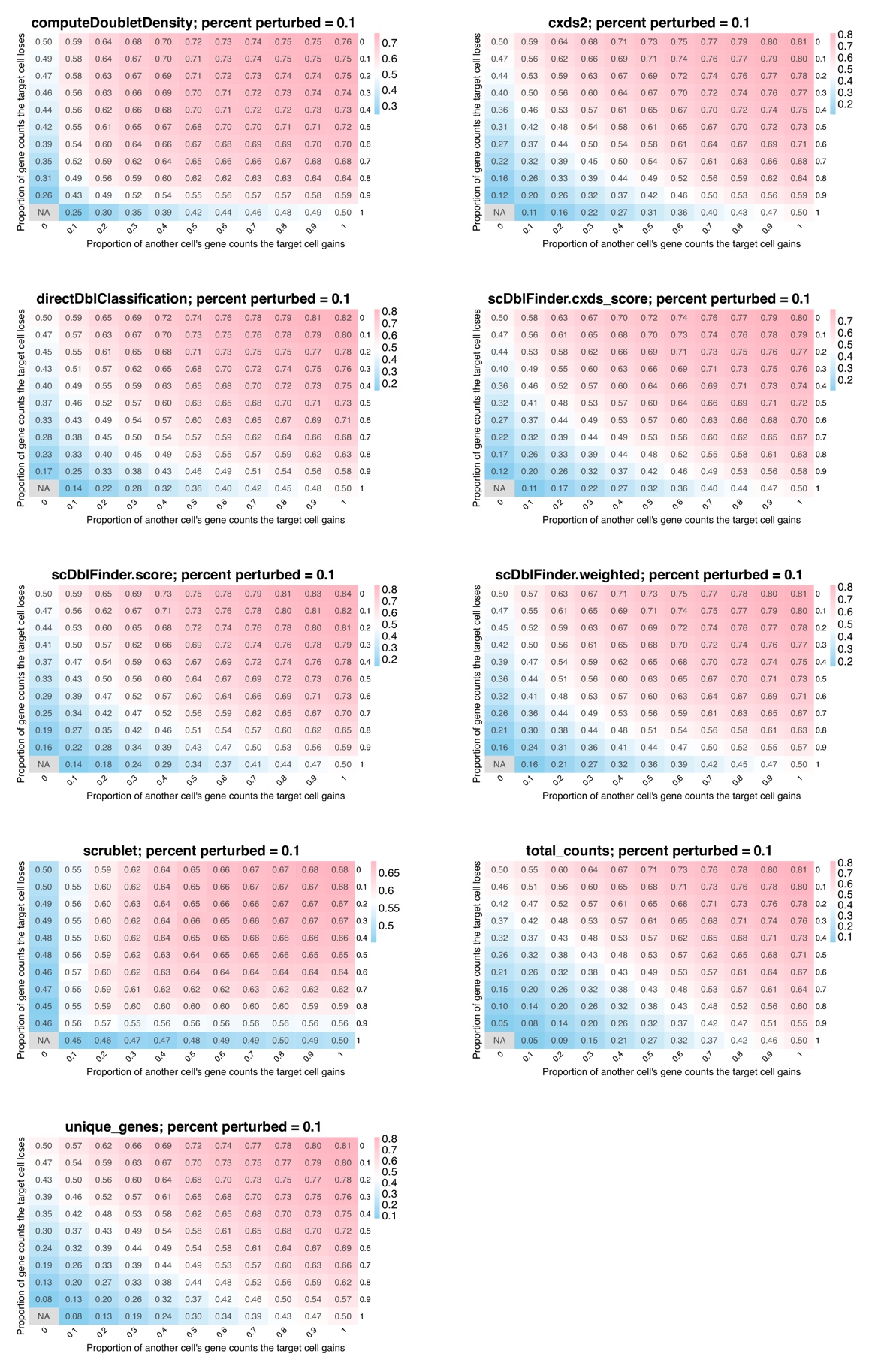

**Figure S14. Error detection in spatial transcriptomic data using 10^th^ neighbor donor cells, with 10% perturbation rate, related to Figure 4.** Heatmaps show AUROCs across the perturbation space for all methods. Each entry shows AUROC averaged over 16 gene panels. In each case, 10% of cells were perturbed. Created in Biorender.com.

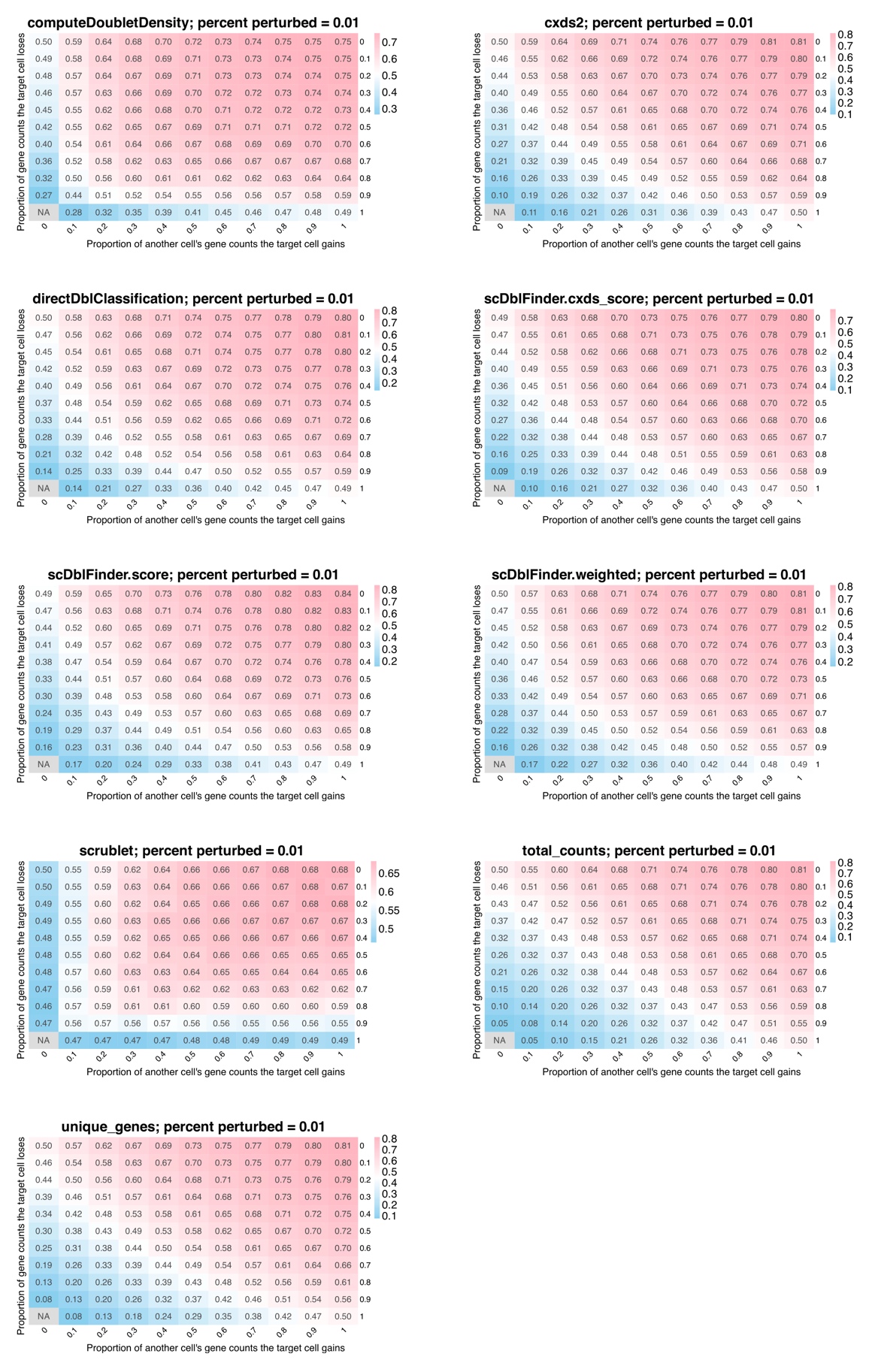

**Figure S15. Error detection in spatial transcriptomic data using 10^th^ neighbor donor cells, with 1% perturbation rate, related to Figure 4.** Heatmaps show AUROCs across the perturbation space for all methods. Each entry shows AUROC averaged over 16 gene panels. In each case, 1% of cells were perturbed. Created in Biorender.com.

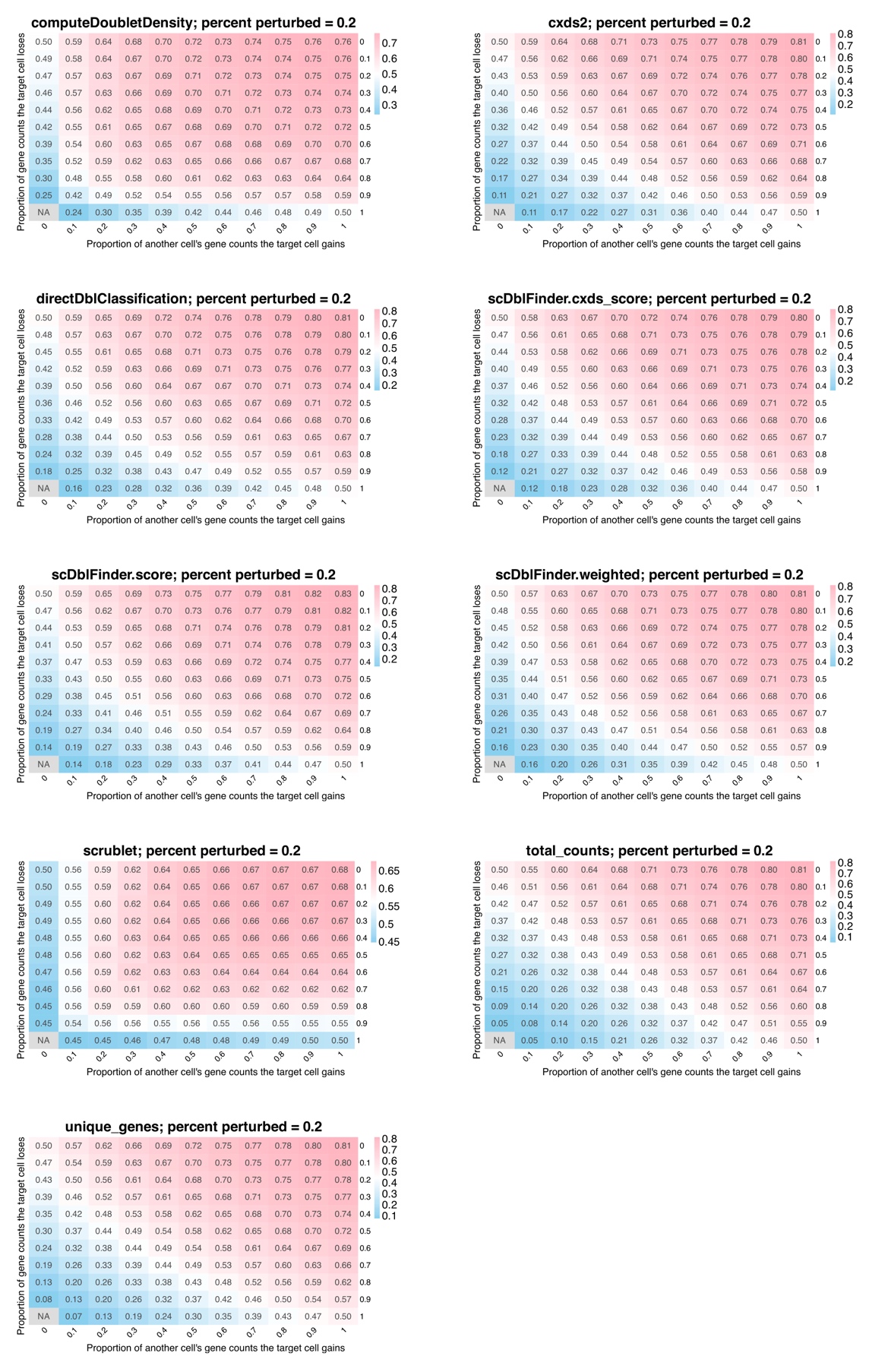

**Figure S16. Error detection in spatial transcriptomic data using 10^th^ neighbor donor cells, with 20% perturbation rate, related to Figure 4.** Heatmaps show AUROCs across the perturbation space for all methods. Each entry shows AUROC averaged over 16 gene panels. In each case, 20% of cells were perturbed. Created in Biorender.com.
